# DNMT1-Mediated Epigenetic Reprogramming Drives *KMT2A* Amplifications and Rearrangements

**DOI:** 10.64898/2026.09.21.753283

**Authors:** Madison A. Lineberger, Zach H. Gray, Benjamin I. Ferman, Erin O’Donnell, Rebecca G. Smith, Kathleen Schiela, Anushka Udara, Kashish Chetal, Chloe Azadegan, Monika Maria Toma, Ethan Sumner, John Santoro, Christopher Miranda, Manna Ahmed, Jade Wilson, William Lautert-Dutra, Chencheng Li, Aidan McKelvey, H. Ümit Kaniskan, Wenyun Zhang, Jian Jin, Fabio Puddu, Mark Consugar, Martin Walsh, Alfonso Bellacosa, Cihangir Duy, Hayan Lee, Tomasz Skorski, Yu Liu, Ruslan I. Sadreyev, Johnathan R. Whetstine

## Abstract

*MLL/KMT2A* amplifications and rearrangements are prevalent in infant, adult, and therapy-induced leukemia; however, the molecular contributors controlling these alterations remain elusive. Here, we reveal that rapid CTCF degradation generates local genome structure alterations at *KMT2A* and copy gains and rearrangements. We then established a conserved, coordinated interplay between DNA and histone methylation pathways that control CTCF occupancy, and in turn, *KMT2A* locus stability. For example, DNMT1 overexpression promotes increased H3K9 methylation, reduced CTCF occupancy, and causes *KMT2A* alterations. However, DNMT1 inhibition suppresses these events and *KMT2A* alterations caused by topoisomerase II inhibition. Locus-specific epigenome targeting demonstrated that DNA or H3K9 methylation promotes *KMT2A* copy gains and rearrangement, whereas targeted TET activity suppresses these events upon methylation perturbation or doxorubicin treatment. These findings identify a conserved, coordinated DNA–histone methylation axis governing *KMT2A* amplifications and rearrangement susceptibility, revealing biomarkers and therapeutic targets to predict and intercept these events in cancer.

**STATEMENT OF SIGNIFICANCE:** This study reveals a conserved DNA-histone methylation mechanism controlling the susceptibility of the *KMT2A* locus to copy gain and rearrange. These events can be generated or prevented by genetic, therapeutic or epigenome editing. Furthermore, DNMT1 inhibition blocks chemotherapy-induced *KMT2A* disruption, identifying a therapeutic strategy to intercept these oncogenic genomic alterations.

## INTRODUCTION

The *KMT2A/MLL* locus undergoes amplification and rearrangement in acute myeloid leukemia (AML) and myelodysplastic syndrome (MDS), where these alterations are associated with poor clinical outcome [1–6]. *KMT2A* alterations are also recurrent in therapy-related leukemias arising after exposure to topoisomerase II (Topo II) inhibitors [7–11]. Despite their clinical significance, the mechanisms that render *KMT2A* susceptible to genomic alteration, particularly following therapeutic stress, remain incompletely understood.

Disruption of chromatin architectural regulators such as CCCTC-binding factor (CTCF) perturbs genome topology, promotes replication stress and defective DNA damage repair, and contributes to chromosomal instability, a hallmark of cancer [12–15]. Consistent with this relationship, alterations in chromatin architecture are associated with copy-number changes and genomic rearrangements across human cancers [16–21]. We previously demonstrated that loss or inhibition of the H3K9me1/2 demethylase KDM3B, or increased activity of the H3K9me1/2 methyltransferase G9a/EHMT2, promotes site-specific *MLL/KMT2A* amplification and rearrangement across hematologic and non-hematologic cell models [17, 21]. These alterations were accompanied by loss of CTCF occupancy within the *KMT2A* breakpoint cluster region (BCR) [17]. Moreover, the Topo II inhibitor doxorubicin reduced KDM3B and CTCF protein levels and induced *KMT2A* alterations, which were suppressed by genetic or pharmacologic inhibition of G9a *in vitro* and *in vivo* [17]. A key question remained about how KDM3B- and G9a-dependent H3K9 methylation is actually coupled to the broader CTCF occupancy landscape at *KMT2A*. Furthermore, the impact of acute CTCF loss on the local *KMT2A* genome architecture and associated alterations remained unclear.

CTCF occupancy is sensitive to local and regional DNA methylation [22–24], providing a potential mechanistic link between H3K9 methylation and CTCF regulation. G9a functionally interacts with DNMT1 to coordinate H3K9 and DNA methylation, including during DNA replication [25, 26]. DNMT1 is also highly expressed in AML and contributes to leukemic maintenance [27, 28]. Disruption of ten-eleven translocation (TET) dependent DNA demethylation is also a recurrent feature of myeloid malignancies, where TET2 loss or mutant IDH1/2-mediated inhibition of TET activity produces aberrant DNA methylation and impairs hematopoietic differentiation [29–31]. These observations led us to hypothesize that coordinated regulation of histone and DNA methylation controls CTCF occupancy across the *KMT2A* region and thereby determines susceptibility to amplification and rearrangement.

Here, we demonstrate that acute CTCF degradation disrupts occupancy at multiple CTCF sites across the *KMT2A*-associated region, perturbs local three-dimensional genome organization, and is sufficient to promote *KMT2A* amplification and rearrangement. Loss of KDM3B alters both 5-methylcytosine (5mC) and 5-hydroxymethylcytosine (5hmC) across the locus, and DNMT1-dependent methylation contributes to regulation of the broader CTCF-occupancy landscape and promotes *KMT2A* genomic alterations. Conversely, DNMT1 depletion or inhibition reduces KDM3B-dependent chromatin changes and suppresses *KMT2A* alterations, whereas DNMT1 overexpression is sufficient to increase H3K9me1, reduce CTCF occupancy, and promote amplification and rearrangement. Inhibition of TET enzymes induces *KMT2A* alterations in a DNMT1- and G9a-dependent manner, demonstrating that active DNA demethylation limits the frequency of *KMT2A* amplification and rearrangement events. Locus-specific epigenome editing further provides functional evidence for this relationship: targeted recruitment of the catalytic domains of DNMT3A or G9a to the *KMT2A* breakpoint cluster region (BCR) is sufficient to induce genomic alterations, whereas targeted TET catalytic activity suppresses alterations driven by epigenetic disruption or Topo II inhibition. Finally, genetic or pharmacologic inhibition of DNMT1 prevents doxorubicin-induced *KMT2A* alterations in human cells and *in vivo*. Together, these findings define a coordinated DNA- and histone-methylation pathway that regulates the CTCF-occupied chromatin domain at *KMT2A* and contributes to susceptibility to genomic alterations. These studies further identify DNMT1-dependent methylation as a potential point of interception for limiting therapy-induced *KMT2A* amplification and rearrangement events.

## RESULTS

### Acute CTCF loss impacts *KMT2A* alterations and genome structure

Disruption of CTCF occupancy has been associated with *MLL/KMT2A* amplification and rearrangement in multiple cell types [17]. However, whether acute CTCF loss is sufficient to rapidly induce these alterations, and how CTCF depletion affects three-dimensional chromatin organization at the *KMT2A* locus, remained unclear. To directly test the effect of acute CTCF loss on *KMT2A*, we used HCT116 cells harboring an auxin-inducible CTCF degron, which enables rapid and reversible depletion of CTCF. We are able to assess the conserved role of CTCF at *KMT2A* and define the immediate consequences of CTCF loss versus prolonged depletion [32]. Public CTCF ChIP-seq data from auxin-inducible HCT116 cells demonstrated prominent occupancy within the *KMT2A* BCR, including exon 11, that was reduced following CTCF degradation **(Fig. 1A, top)** [33]. Following 24 hours of adherence, CTCF degradation was achieved by treatment with the synthetic auxin analog 5-phenyl-indole-3-acetic acid (5-Ph-IAA) at 1 μM for 1 hour without altering KDM3B protein levels (**Fig. 1A, bottom**), allowing us to directly assess the immediate consequences of CTCF loss independently of changes in KDM3B.

**Fig. 1.**
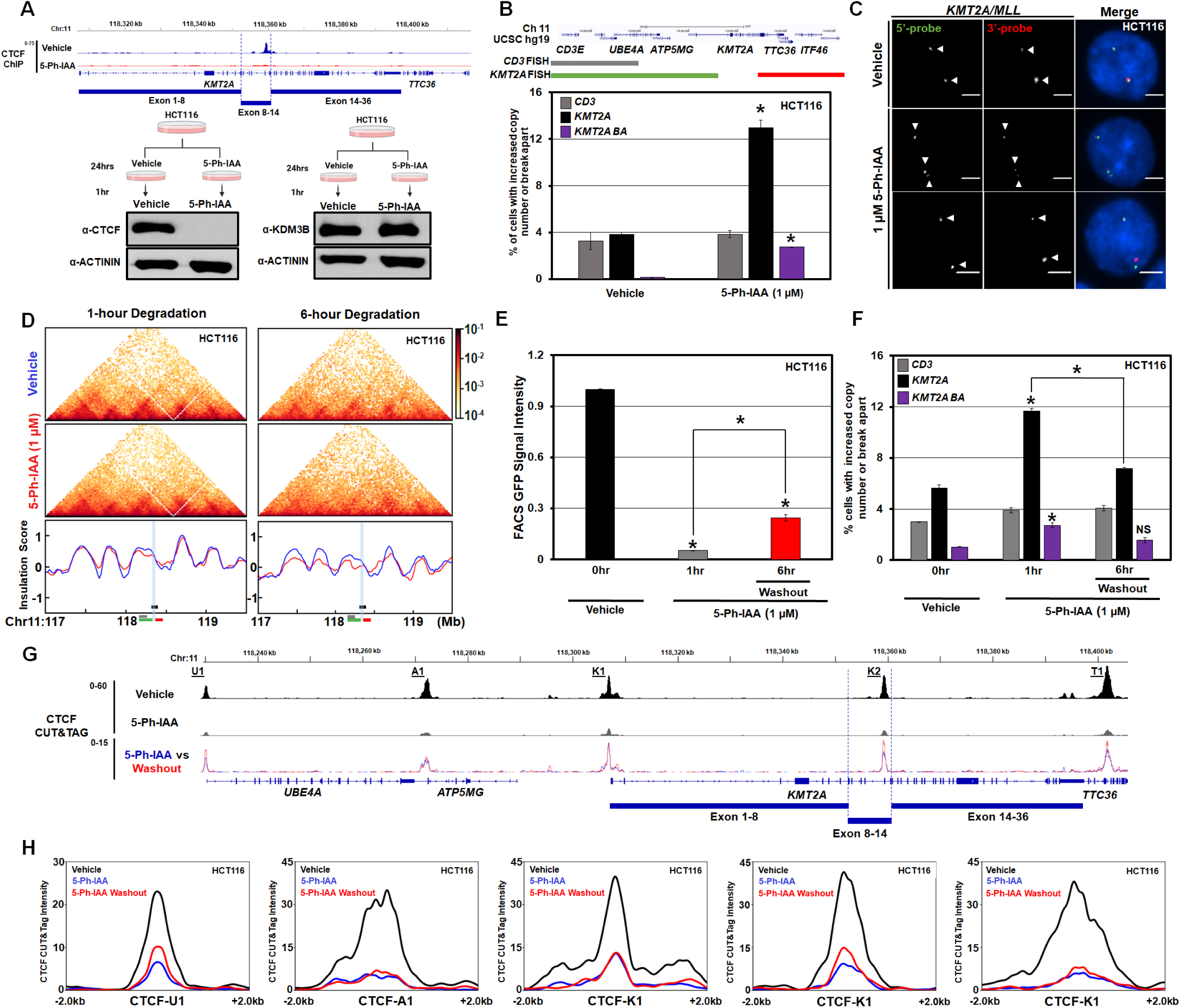
Rapid loss of CTCF drives *KMT2A/MLL* genomic alterations. (A) Public CTCF ChIP-seq in HCT116 Schematic showing depletion of CTCF at *KMT2A* (upper; [33]). Schematic of the HCT-116 degron system. Representative western blots demonstrate CTCF protein levels are reduced within one hour of 1 µM 5-Ph-IAA treatment (left), while KDM3B protein levels are not impacted (right). (B) Upper schematic shows the DNA fluorescent in situ hybridization (FISH) probe genomic locations that are used for *KMT2A* locus. Treatment with 1 µM 5-Ph-IAA in one hour generate *KMT2A* copy gains (black) and break aparts (purple) with no alterations to the adjacent control *CD3* region (grey). (C) Representative images with the *KMT2A* DNA FISH break apart probe (red and green probes in panel B) that show no copy gain in Vehicle (top panel), DNA copy gains (middle panel) and break apart events (lower panel) upon 5-Ph-IAA treatment. Arrowheads highlight the FISH signal. (D) Hi-C analysis in HCT116 cells demonstrating rapid degradation of CTCF via Auxin-inducible system alters 3D genome interactions at the *KMT2A* locus within 1 hr (left) and 6 hrs (right). *KMT2A* and the DNA FISH probes in Fig. 1B are noted. The BCR is highlighted in blue shading. (E) Re-expression of CTCF quantified after depletion by the Auxin-inducible system via GFP-FACS. HCT116 cells were treated with 1 µM 5-Ph-IAA for 1 hr before medium exchange. Samples were taken 6 hrs later for FACS analysis. (F) DNA FISH of HCT116 cells treated with 1 µM 5-Ph-IAA for 1 hr and collected 6 hrs after drug washout. (G) CTCF CUT&Tag data at the *KMT2A* locus in HCT116 cells (black) demonstrating reduced CTCF occupancy following 1 hr of 1 µM 5-Ph-IAA treatment (gray) and partial restoration 6 hrs after drug washout (red overlay with blue). Example CTCF peaks are noted with first letter of nearest gene. Multiple CTCF peaks within one gene are numbered accordingly. (H) CTCF CUT&Tag signal profiles centered on CTCF sites spanning the *KMT2A* region in HCT116 cells treated with vehicle (black), 5-Ph-IAA (gray), or following 1 µM 5-Ph-IAA washout (red) (CTCF-U1 to CTCF-T1; panel G). Normalized CTCF CUT&Tag signal is shown across ±2 kb surrounding each site. Profiles represent mean spike-in-normalized signal across biological replicates. Error bars represent the SEM. Asterisk indicates significant difference from indicated (p < 0.05) by two-tailed Student’s t test. A minimum of 2 replicates per experiment were conducted. NS, not significant to control. Scale bar represents 5 μm. Representative DNA FISH images were adjusted for figure presentation only-see materials and methods.

We next determined whether acute CTCF depletion was sufficient to induce *KMT2A* genomic alterations. DNA FISH was used to assess the locus. Probes targeting the 5′ and 3′ regions of *KMT2A* were used to distinguish focal copy-number gains from break-apart events, whereas an adjacent, partially overlapping control probe (CD3) was used to assess the focality of the alterations (**Fig. 1B)** [17]. Copy number and break apart events were quantified per nucleus as previously described [17, 34–38]. Experimental conditions were independently validated, and flow-cytometric analysis demonstrated no significant change in cell-cycle distribution following 1 hour of CTCF degradation (**Supplemental Fig. S1A**). Despite this short exposure, acute CTCF loss resulted in a significant increase in *KMT2A* amplification and rearrangement events (**Fig. 1B**). Representative DNA FISH images show two paired 5′ and 3′ regions of the *KMT2A* break-apart probe in vehicle-treated cells (**Fig. 1C, upper**). In contrast, 5-Ph-IAA treated cells show focal amplification (**Fig. 1C, middle**) and/or separation of the 5’ and 3’ probe signals consistent with *KMT2A* rearrangement (**Fig. 1C, lower**). These findings demonstrate that acute CTCF depletion is sufficient to rapidly increase *KMT2A* genomic alterations in the absence of detectable cell-cycle changes.

We next asked whether these genomic events were accompanied by changes in local three-dimensional chromatin organization. Hi-C contact maps showed altered chromatin interaction patterns surrounding *KMT2A* within 1 hour of CTCF degradation (**Fig. 1D, left; Supplemental Fig. S1B**). After 6 hours, these alterations extended across a broader region flanking the *KMT2A* locus (**Fig. 1D, right; Supplemental Fig. S1B**). Consistent with these findings, publicly available Hi-C data from HAP1 cells showed similar alterations in chromatin interaction patterns surrounding *KMT2A* following 48 hours of CTCF degradation, highlighting the conserved role of CTCF in modulating the *KMT2A* locus and preventing these genomic alterations (**Supplemental Fig. S1B**) [39]. These data demonstrate that loss of CTCF rapidly perturbs three-dimensional chromatin organization across the broader *KMT2A* domain that coincides with the focal copy gains and rearrangement.

Because CTCF depletion was reversible in this system, we determined whether restoration of CTCF could re-establish occupancy at *KMT2A* and suppress the genomic phenotype. Following 5-Ph-IAA washout, CTCF-GFP signal significantly recovered within 6 hours, demonstrating partial restoration of CTCF protein levels (**Fig. 1E**), which was accompanied by a complete rescue of *KMT2A* copy gains and rearrangements (**Fig. 1F**). Furthermore, CTCF CUT&Tag demonstrated substantial loss of occupancy at regional CTCF sites following 1 hour of degradation (**Fig. 1G-H**, **Supplemental Fig. S1C**). Following washout, CTCF occupancy showed partial recovery across multiple sites spanning the *KMT2A* region, including the BCR (CTCF-4) and the proximal and distal sites to *KMT2A.* The degree of recovery varied among sites (**Fig. 1G-H**). Consistent with this partial recovery, comparative CUT&Tag analysis after washout showed a higher concordance of CTCF signal with the vehicle state, supporting partial restoration of CTCF occupancy across the locus (**Supplemental Fig. S1C**). ChIP-qPCR independently confirmed increased CTCF occupancy at the *KMT2A* BCR by 24 hours following washout (**Supplemental Fig. S1D**).

These findings demonstrate that acute CTCF loss is associated with reduced CTCF occupancy across the *KMT2A* region, coincident with disruption of local chromatin interactions and increased *KMT2A* amplification and rearrangement. Partial recovery of CTCF is accompanied by increased occupancy at multiple CTCF sites across the region, and in turn, suppression of *KMT2A* genomic alterations. These findings highlight the regional plasticity and support a model in which the broader CTCF-occupied domain surrounding *KMT2A* contributes to maintenance of local architecture and the inhibition of copy gains and rearrangements.

### KDM3B and DNMT1 coordinate the *KMT2A* chromatin landscape to regulate CTCF occupancy and genomic alterations

CTCF binding is sensitive to CpG methylation within its recognition motif and surrounding sequences [22–24] and inhibition of DNA demethylation can reduce CTCF occupancy [22, 29]. DNA and histone methylation are also coordinated through interactions between DNMTs and the H3K9 methyltransferase G9a/EHMT2 [25, 26]. We previously demonstrated that the H3K9me1/2 demethylase KDM3B regulates H3K9 methylation at the *KMT2A* locus [17]. Consistent with this relationship, H3K9me1 and H3K9me2 are altered following single, and combined, depletion of KDM3B and G9a/EHMT2, with prominent changes across the *KMT2A* locus (**Fig. 2A**; [17]). Both KDM3B and CTCF occupy multiple regions across the *KMT2A* locus, with prominent enrichment throughout the breakpoint cluster region (BCR) (**Fig. 1G** and **2A**; [17]). CTCF occupancy was reduced upon KDM3B depletion across the *KMT2A*-associated locus (**Fig. 2A**; [17]). Given the established coordination between DNA methylation and H3K9 methylation, these findings raised the possibility that perturbation of KDM3B-dependent H3K9 methylation may also alter the DNA methylation state of *KMT2A*, thereby affecting CTCF occupancy. We therefore hypothesized that DNMT-dependent DNA methylation contributes to the epigenetic regulation of *KMT2A* and influences the generation of genomic alterations at this locus (**Supplemental Fig. S2A**).

**Fig. 2.**
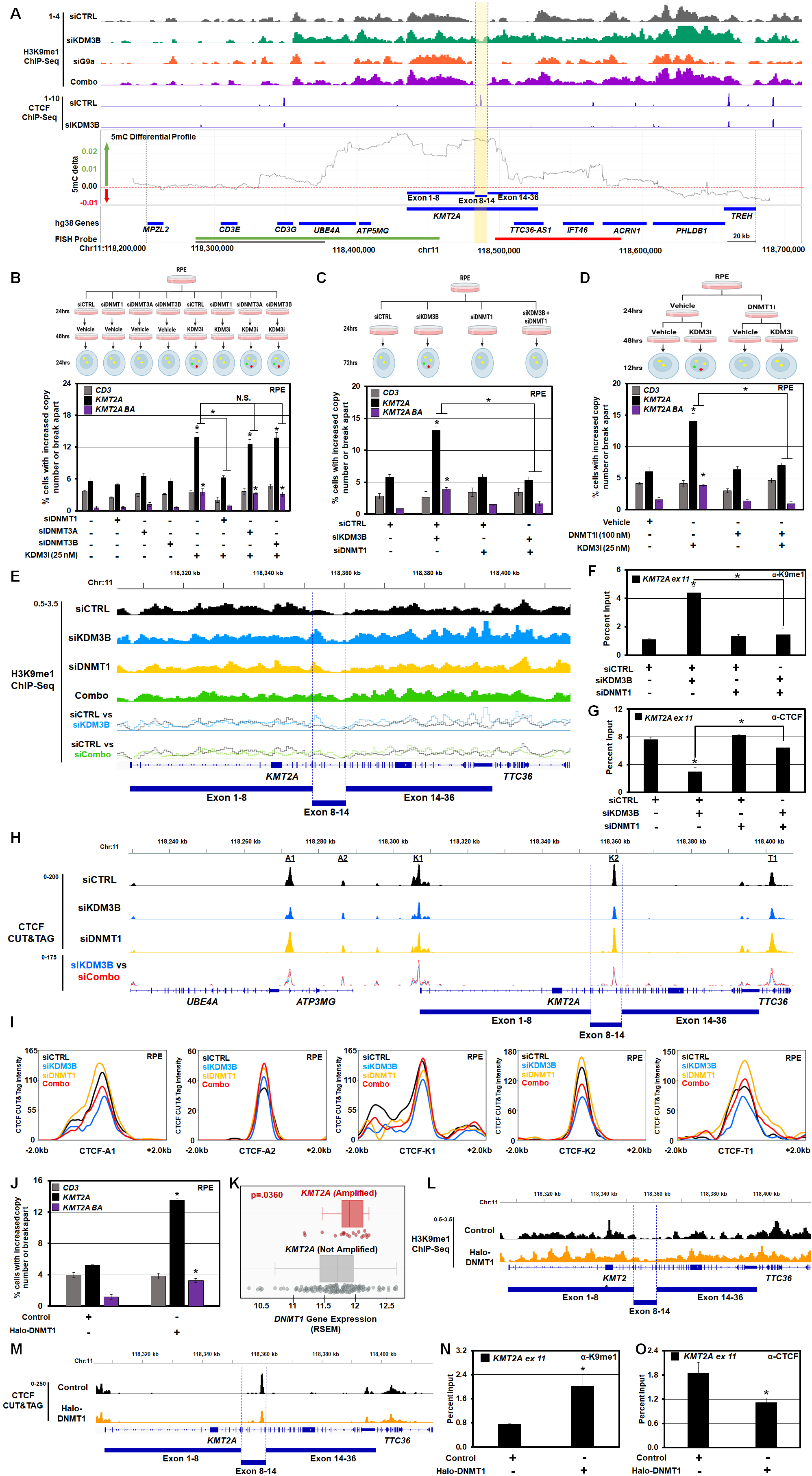
DNMT1 controls *KMT2A* genomic alterations caused by KDM3 perturbation. (A) Publicly available H3K9me1 ChIP-seq for siKDM3B, siG9a, and the combination (top; [17]). Publicly available CTCF ChIP-seq data in siKDM3B-depleted cells (middle; [17]). Differential methylation between control and KDM3B siRNA knockdown is shown as rolling averages for a replicate (bottom panel). The green arrow notes positive 5mC delta versus the negative delta with the red arrow. Bottom panel labels include gene model derived from the UCSC Known Genes track. Red, green, and grey lines indicate the positions of the corresponding DNA FISH probes. The yellow band indicates exons 8–14 of *KMT2A*, which coincides with the BCR. (B) Schematic (upper) and DNA FISH (lower) demonstrating that DNMT1 siRNA depletion rescues KDM3i-induced *KMT2A* alterations. (C) Schematic (upper) and DNA FISH (lower) demonstrating depletion of KDM3B and DNMT1 rescues *KMT2A* alterations caused by KDM3B depletion alone. (D) Schematic (upper) and DNA FISH (lower) demonstrating that KDM3i treatment promotes *KMT2A* alterations, but 100 nM DNMT1i co-treatment rescues *KMT2A* events in RPE cells. 100 nM DNMT1i alone has no impact on the *KMT2A* locus. (E) ChIP-seq for H3K9me1 in RPE cells treated with siControl (black), siKDM3B (blue), siDNMT1 (yellow), or siRNA co-depletion (Combo, green) are shown. Line plots for siControl (siCTRL, black) compared to siKDM3B (blue) or to siKDM3B/siDNMT1 (Combo, green) are shown to illustrate the reduced H3K9me1 observed when DNMT1 is co-depleted with KDM3B. (F) ChIP-qPCR demonstrating increased H3K9me1 at *KMT2A* exon 11 following KDM3B siRNA depletion. siRNA co-depletion with DNMT1 prevents the increased H3K9me1. (G) ChIP-qPCR demonstrating decreased CTCF occupancy at *KMT2A* exon 11 following KDM3B depletion that is partially restored following DNMT1 co-depletion. (H) CTCF CUT&Tag tracks are shown at the *KMT2A* locus, which illustrates reduced CTCF occupancy when KDM3B is depleted and partial recovery upon co-depletion with DNMT1. Example CTCF called peaks are noted with first letter of nearest gene. Multiple CTCF peaks within one gene are numbered accordingly. (I) CTCF CUT&Tag signal profiles centered on CTCF sites (CTCF-A1–CTCF-T1) spanning the *KMT2A* region following siControl (black), siKDM3B (blue), siDNMT1 (yellow), or siRNA co depletion (Combo, red) depletion in RPE cells. Normalized CTCF CUT&Tag signal is shown across ±2 kb surrounding each site. Profiles represent mean signal across biological replicates. (J) DNA FISH demonstrating DNMT1 overexpression promotes *KMT2A* events in RPE cells. (K) Analysis of TCGA LAML samples with *KMT2A* amplifications. Plot shows that most LAML samples that are *KMT2A* amplified also have increased DNMT1 expression (RSEM) with a p-value of 0.0360. Statistical significance was computed by Wilcoxon rank-sum test, which provides a non-parametric hypothesis test on two independent samples. (L) ChIP-seq for H3K9me1 in DNMT1 overexpressed RPE cells. (M) CTCF CUT&Tag tracks at the *KMT2A* locus demonstrating reduced CTCF occupancy upon DNMT1 overexpression in RPE cells. (N) ChIP-qPCR demonstrating an increase of K9me1 at *KMT2A* exon 11 (*KMT2A* ex 11) following DNMT1 overexpression in RPE cells. (O) ChIP-qPCR demonstrating reduced CTCF occupancy at *KMT2A* exon 11 (*KMT2A* ex 11; black) following DNMT1 overexpression in RPE cells. Error bars represent the SEM. Asterisk indicates significant difference from indicated (p < 0.05) by two-tailed Student’s t test. A minimum of 2 replicates per experiment were conducted.

To determine whether KDM3B regulates DNA methylation at *KMT2A*, we performed genome-wide 5-methylcytosine (5mC) profiling using the biomodal duet evoC assay following KDM3B depletion via siRNA in the retinal-pigment epithelial (RPE-1; called RPE) cells (**Fig. 2A**). These cells were used because of their near-diploid, genomically stable background and their established utility for studying the conserved amplification and rearrangement mechanisms [17, 34, 37, 40–44]. In fact, RPE cells were used to discover the conserved regulation of *KMT2A* amplification and rearrangement control [17]. KDM3B depletion increased 5mC across the *KMT2A* locus in biological replicates (**Fig. 2A; Supplemental Fig. S2B**), whereas methylation changes at other genomic regions were less consistent between replicates. These data suggest that KDM3B restrains 5mC accumulation at the *KMT2A* locus and suggest that loss of KDM3B alters both the DNA and histone methylation environment surrounding the *KMT2A* BCR.

Given the increase in 5mC following KDM3B depletion, we next determined which DNMT enzyme(s) were required for KDM3 enzymatic inhibitor-induced (KDM3i) *KMT2A* genomic alterations [17]. DNMT1, DNMT3A, and DNMT3B were individually depleted using two independent siRNAs per gene in RPE cells treated with KDM3i (**Fig. 2B**). Depletion with each siRNA was validated prior to analysis (**Supplemental Fig. S2C**). Among the DNMT family members tested, only DNMT1 depletion suppressed KDM3i-induced *KMT2A* genomic alterations, whereas depletion of DNMT3A or DNMT3B was insufficient to prevent these events (**Fig. 2B**). This relationship was confirmed genetically, as DNMT1 depletion rescued the increase in *KMT2A* amplification and rearrangement caused by KDM3B siRNA depletion (**Fig. 2C; Supplemental Fig. S2D-E**). Consistent with this rescue, DNMT1 depletion also reduced 5mC at the *KMT2A* locus (**Supplemental Fig. S2F-G**). Together, these data identify DNMT1 as the DNA methyltransferase required for DNA methylation and the *KMT2A* genomic alterations following KDM3B depletion or inhibition.

We next asked whether pharmacologic inhibition of DNMT1 was sufficient to prevent KDM3i-induced *KMT2A* alterations. Cells were treated with the selective DNMT1 inhibitor GSK-3685032 (DNMT1i; 100 nM; [45]) prior to KDM3i treatment. DNMT1 inhibition suppressed KDM3i-induced *KMT2A* alterations without affecting cell growth or cell-cycle distribution (**Fig. 2D; Supplemental Fig. S2H-I**). DNMT1 inhibition also had a conserved role in blocking KDM3i-induced *KMT2A* copy gains and rearrangements in an AML-derived cellular model (**Fig. S2J**). Treatment with the DNA methyltransferase inhibitor 5-azacytidine (5-AzaC), an FDA-approved epigenetic therapeutic agent [46, 47], also prevented KDM3i-induced *KMT2A* amplification and rearrangement without significantly altering cell growth or cell-cycle distribution at the concentration examined (**Supplemental Fig. S2K-M**). Thus, both genetic and pharmacologic inhibition of DNMT1 suppress *KMT2A* genomic alterations following KDM3B inhibition, supporting a critical role for DNA methylation in the generation of these alterations.

We then determined whether DNMT1 was impacting the increased H3K9 methylation that was generated by KDM3B depletion. Consistent with a prior report (**Fig. 2A**; [17]), KDM3B loss increased H3K9me1 across the *KMT2A* locus (**Fig. 2E**). The increased H3K9me1 was reduced upon DNMT1 co-depletion*KMT2A* (**Fig. 2E**). Targeted ChIP-qPCR at a representative region within the BCR independently confirmed the reduction in H3K9me1 following combined KDM3B and DNMT1 depletion (**Fig. 2F**). These data demonstrate that DNMT1 contributes to KDM3B-dependent H3K9me1 accumulation across the *KMT2A* locus.

Because DNA methylation and H3K9 methylation can influence CTCF binding, we next determined whether DNMT1 contributes to the altered CTCF occupancy landscape produced by KDM3B loss. Targeted ChIP-qPCR confirmed reduced CTCF occupancy within the *KMT2A* BCR following KDM3B depletion and partial restoration upon co-depletion of DNMT1 (**Fig. 2G**). CTCF CUT&Tag further demonstrated that KDM3B depletion reduced occupancy at multiple CTCF-bound regions spanning the *KMT2A* locus, rather than selectively affecting a single site, with the magnitude of loss varying across the region **(Fig. 2H, I).** Co-depletion of KDM3B and DNMT1 partially restored CTCF occupancy at multiple affected regions (**Fig. 2I**), which was similar to the partial rescue observed upon washout in the CTCF-degron cells (**Fig. 1H**). Together, these findings indicate that KDM3B loss disrupts CTCF occupancy across the broader *KMT2A*-associated domain, promotes DNMT1-dependent methylation, and that DNMT1 contributes to the site-dependent loss of CTCF binding throughout this region. These data further emphasize a coordinate relationship between H3K9 and DNA methylation at the *KMT2A* locus.

To further test this relationship, we assessed whether increased DNMT1 expression was sufficient to recapitulate the genomic and chromatin changes observed following KDM3B depletion. DNMT1 overexpression was sufficient to increase *KMT2A* copy gains and rearrangements (**Fig. 2J; Supplemental Fig. S2N**). Consistent with this data, increased *DNMT1* expression was significantly higher in *KMT2A*-amplified patient samples compared to non-amplified (**Fig. 2K**). This relationship was not observed for the other DNMTs (**Supplemental Fig. S2O**) H3K9me1 was also increased across the *KMT2A* locus following DNMT1 overexpression, which was similar to the increase with KDM3B depletion (**Fig. 2A, E, L**). Consistent with this relationship, G9a depletion rescued the *KMT2A* alterations caused by DNMT1 overexpression, while DNMT1 depletion suppressed the G9a-induced events (**Supplemental Fig. S2P-S**). Furthermore, CTCF CUT&Tag demonstrated a corresponding reduction in CTCF occupancy across the *KMT2A* locus following DNMT1 overexpression (**Fig. 2M**). Targeted ChIP-qPCR within the BCR independently confirmed increased H3K9me1 and reduced CTCF occupancy (**Fig. 2N-O**). Together, these findings demonstrate that increased DNMT1 expression is sufficient to promote *KMT2A* genomic alterations and reproduce the increased H3K9me1 and reduced CTCF occupancy observed following KDM3B loss. These data further illustrate the coordinated control of this locus by DNMT1 and G9a.

### TET-mediated DNA demethylation controls *KMT2A* copy gains and rearrangements

At methylation-sensitive CTCF-binding sites, increased 5mC can interfere with CTCF occupancy, whereas oxidation of 5mC to 5hmC by TET enzymes represents an intermediate step in active DNA demethylation [31, 47]. Disruption of DNA methylation homeostasis can alter local chromatin organization and compromise CTCF-dependent genome regulation [48]. Given the increase in 5mC following KDM3B depletion and the requirement for DNMT1 in promoting *KMT2A* genomic alterations, we hypothesized that active DNA demethylation counteracts *KMT2A* amplifications and rearrangements by maintaining the epigenetic balance at this locus (**Fig. 3A**).

**Fig. 3.**
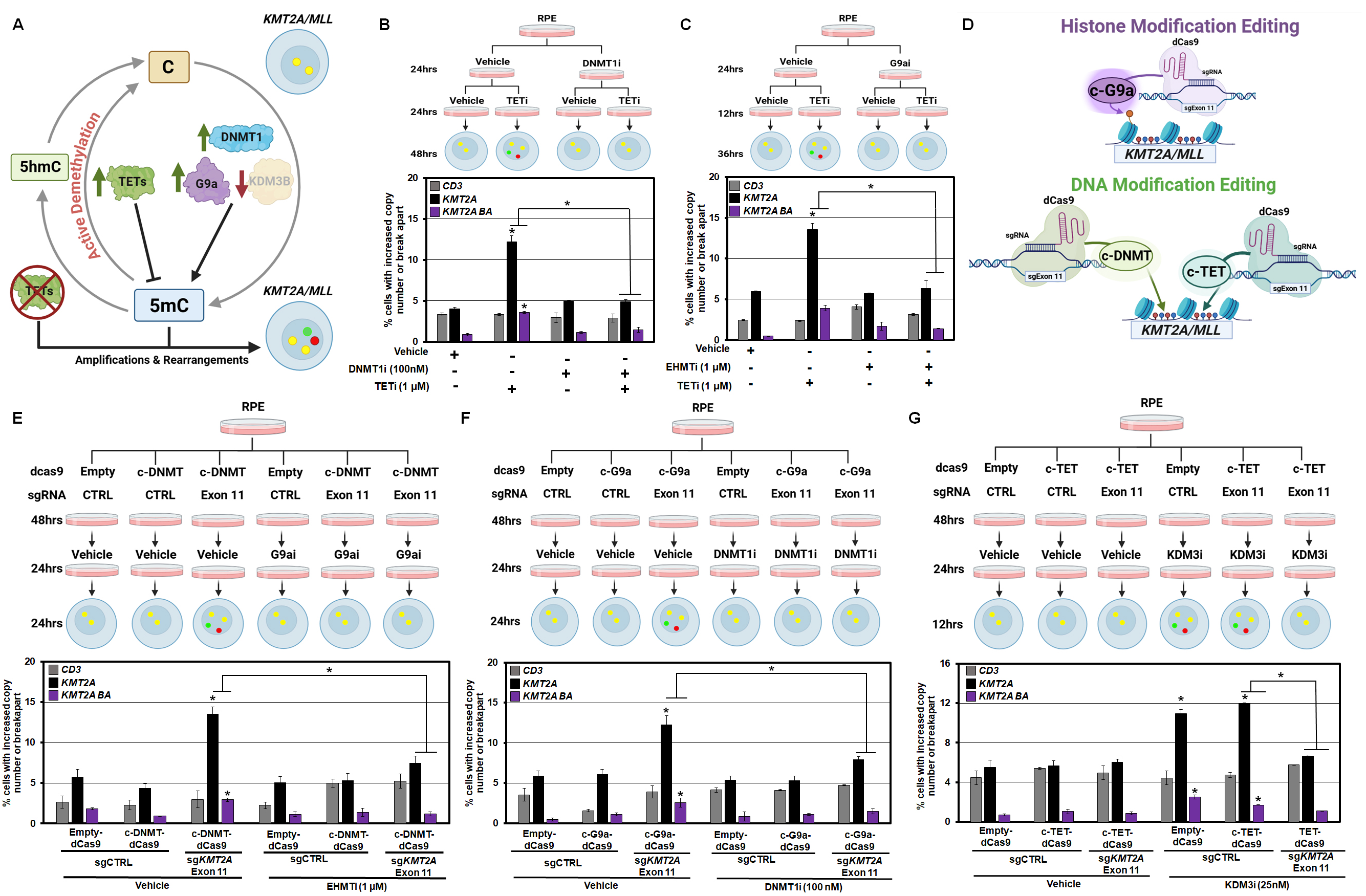
DNA demethylation controls *KMT2A* copy gains and rearrangements. (A) Schematic depicting the DNA methylation cycle and proposed accumulation of 5mC and the amplification and rearrangement phenotypic consequences at the *KMT2A* locus. (B) Schematic (upper) and DNA FISH (lower) demonstrating pre-treatment with DNMT1i prior to 1 µM TETi fully rescues TETi-induced *KMT2A* events. (C) Schematic (upper) and DNA FISH (lower) demonstrating pre-treatment with 1 µM EHMTi prior to 1 µM TETi fully rescues TETi-induced *KMT2A* events. (D) Schematic depicting the DNA and histone methylation c-dCas9 epigenetic editing platform used to evaluate *KMT2A* copy gains and rearrangements. (E) Schematic (upper) and DNA FISH (lower) demonstrating DNMT1i treatment following transfection of c-G9a-dCas9 to exon 11 of *KMT2A* prevents the c-dCas9-induced copy gains and rearrangement. (F) Schematic (upper) DNA FISH (lower) demonstrating that 1 µM EHMTi treatment following transfection of c-DNMT-dCas9 to exon 11 of *KMT2A* prevents the dCas9-induced alterations. (G) Schematic (upper) and DNA FISH (lower) demonstrating that pre-treatment of c-TET-dCas9 to exon 11 of *KMT2A* prior to 25 nM KDM3i fully rescues KDM3i-induced *KMT2A* alterations. Error bars represent the SEM. Asterisk indicates significant difference from indicated (p < 0.05) by two-tailed Student’s t test. A minimum of 2 replicates per experiment were conducted.

Consistent with altered TET-mediated oxidation at this locus, we observed increased 5hmC across *KMT2A* following KDM3B depletion (**Supplemental Fig. S3A-B**). Because 5hmC can represent either a demethylation intermediate or a relatively stable DNA modification, its accumulation alongside 5mC indicates disruption of the balance between DNA methylation and TET-mediated oxidation [31]. The simultaneous accumulation of 5mC and 5hmC suggests an altered methylation equilibrium in which increased DNMT1- and G9a-dependent methylation is accompanied by an increase in TET-mediated oxidation (**Fig. 3A**). We therefore reasoned that inhibition of TET-mediated DNA demethylation should further shift this balance toward methylation and induce *KMT2A* genomic alterations in a DNMT1- and G9a-dependent manner (**Fig. 3A**).

Treatment with the TET inhibitor C35 at 1µM for 72 hours was sufficient to induce *KMT2A* amplification and rearrangement. These alterations were prevented by prior inhibition and genetic depletion of either DNMT1 or G9a (**Fig. 3B-C; Supplemental Fig. S3C-E**). These findings demonstrate that *KMT2A* amplifications and rearrangements induced by TET inhibition depend on both DNMT1 and G9a activity under these conditions. Thus, factors that shift the balance between DNA methylation and active DNA demethylation can influence the frequency of *KMT2A* genomic alterations.

To determine whether altering this epigenetic balance directly at *KMT2A* was sufficient to control genomic alterations, we used dCas9-based epigenome editing constructs containing the catalytic domains of DNMT3a, G9a, or TET1, hereafter referred to as c-DNMT-dCas9, c-G9a-dCas9, and c-TET-dCas9, respectively (**Fig. 3D**) [49, 50]. They were targeted to exon 11 of the *KMT2A* BCR (**Fig. 3D**). Targeted c-DNMT-dCas9 activity at exon 11 was sufficient to induce *KMT2A* genomic alterations, and inhibition of G9a/EHMT activity prevented these events (**Fig. 3E**). Reciprocally, DNMT1 inhibition suppressed *KMT2A* alterations induced by targeted c-G9a-dCas9 activity (**Fig. 3F**). Thus, *KMT2A* alterations induced by targeted DNMT activity depend on G9a activity, and the alterations induced by targeted G9a activity also depend on DNMT1.

We next asked whether targeting TET catalytic activity, specifically at *KMT2A,* could protect the locus from genomic amplifications and rearrangements. Targeting c-TET-dCas9 activity to exon 11 prevented KDM3i-induced *KMT2A* copy gains and rearrangement (**Fig. 3G**). Targeted TET activity also partially suppressed *KMT2A* alterations generated by DNMT1 overexpression (**Supplemental Fig. S3F-G**). Together, these experiments demonstrate that the local DNA methylation state of *KMT2A* is sufficient to influence its susceptibility to genomic alterations. The dependencies among DNMT1, G9a, KDM3B, and TET activity further indicate that the *KMT2A* locus is sensitive to the balance between DNA methylation, active DNA demethylation, and H3K9 methylation state. These data also emphasize the need for the coordinated control by DNMT1-G9a to generate the *KMT2A* copy gains and rearrangements, which highlights a path to suppressing their induced alterations through therapeutic interception.

### DNMT1 Inhibition Suppresses Doxorubicin-Induced KMT*2A* alterations *in vitro* and *in vivo*

Topoisomerase II inhibitors, including doxorubicin (Dox), directly promote *KMT2A* amplification and rearrangement and are associated with reduced KDM3B and CTCF protein levels and decreased CTCF occupancy at *KMT2A* [17]. Enzymatic inhibition of G9a prevents Dox-induced *KMT2A* alterations [17]. Given the functional interaction between DNMT1 and G9a identified above, we tested whether disrupting DNA methylation could similarly suppress Topo II inhibitor-induced *KMT2A* genomic alterations.

DNMT1 depletion via siRNA in RPE cells significantly reduced Dox-induced *KMT2A* copy gains and rearrangement (**Fig. 4A; Supplemental Fig. S4A**). We next tested whether pharmacologic DNMT1 inhibition reproduced this effect. Treatment with the selective DNMT1 inhibitor prevented Dox-induced *KMT2A* genomic alterations in RPE cells and in a primary AML cell line (**Fig. 4B-C; Supplemental Fig. S4B**). These conserved results suggest that DNMT1 activity is required for the generation of *KMT2A* alterations following Dox exposure. We next determined whether targeting TET catalytic activity at *KMT2A* could similarly protect the locus from Dox-induced genomic alterations. Targeting c-TET-dCas9 activity to *KMT2A* exon 11 was sufficient to prevent Dox-induced amplification and rearrangement (**Fig. 4D**). Together with the DNMT1 inhibition studies, these findings demonstrate that the local balance between DNA methylation and active demethylation regulates the susceptibility of *KMT2A* to Topo II inhibitor-induced genomic alterations.

**Fig. 4.**
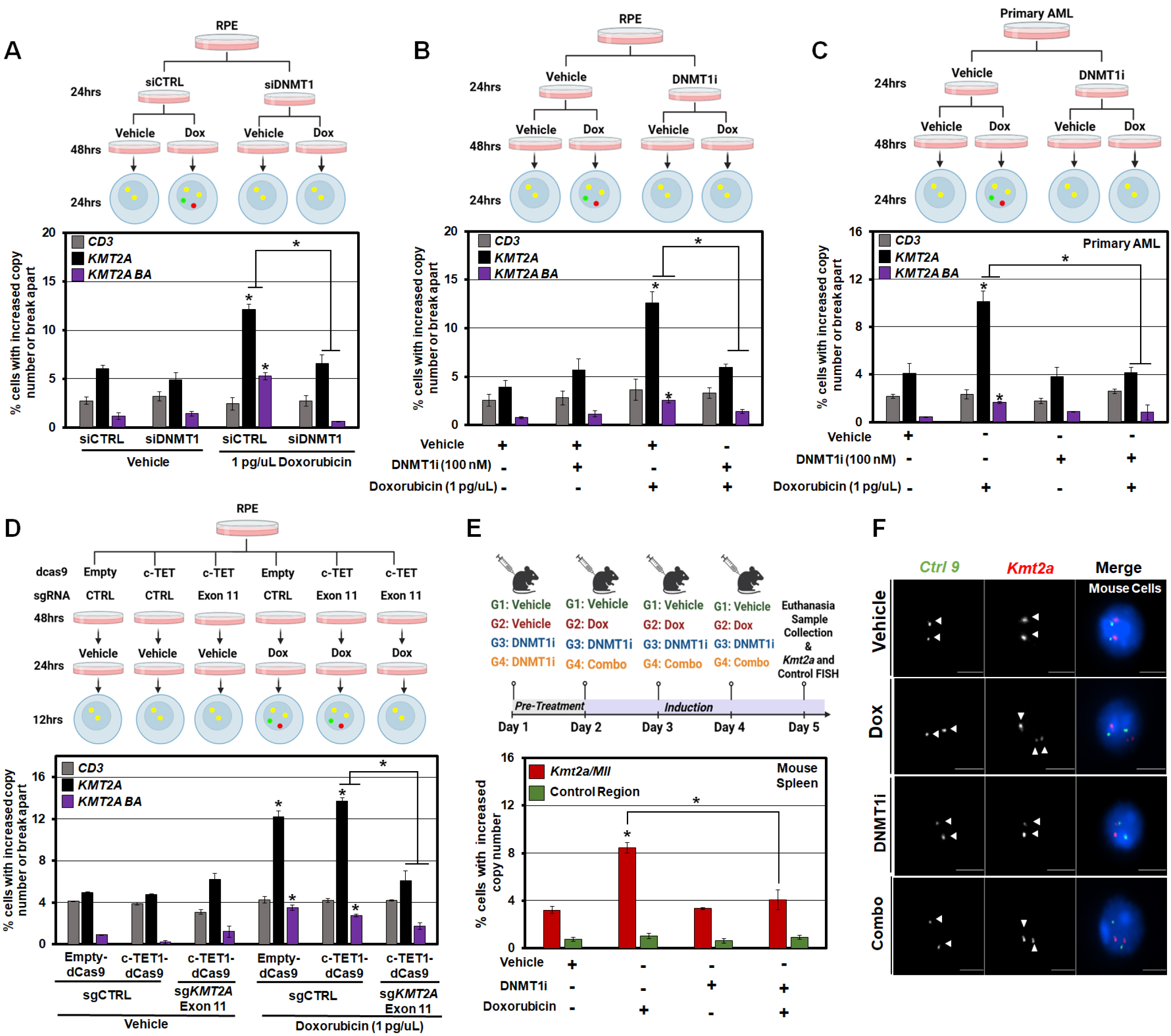
Blocking DNA methylation prevents Topoisomerase II inhibitor-induced *KMT2A/MLL* genomic alterations *in vitro* and *in vivo*. (A) Schematic (upper) and DNA FISH (lower) demonstrate DNMT1 depletion prior to Dox treatment rescues the *KMT2A*-induced alterations in RPE cells. (B) Schematic (upper) and DNA FISH (lower) demonstrate 100 nM DNMT1i prior to 1 pg/uL Dox treatment rescues *KMT2A-*induced genomic alterations in RPE cells. (C) Schematic (upper) and DNA FISH (lower) demonstrate 100 nM DNMT1i prior to 1 pg/uL Dox treatment rescues *KMT2A*-induced alterations in primary AML cells. (D) Schematic (upper) and DNA FISH (lower) demonstrate c-TET-dCas9 directly targeted to exon 11 of *KMT2A* prior to Dox treatment fully rescues the *KMT2A*-induced events in RPE cells. (E) DNMT1i *in vivo* experiment schematic (upper). Pre-treatment of 45mg/kg DNMT1i prior to 1.5mg/kg Dox treatment rescues *Kmt2a/Mll* (red) copy gains *in vivo* (lower). (F) Representative images with *KMT2A/Mll* and control mouse DNA FISH probes (red and green, respectively) that show no copy gains in Vehicle treated (top panel), DNA copy gains with Dox treatment (second row of panels), and no DNA copy gains in DNMT1i alone (third row) or mice receiving combination treatment (fourth row). Arrowheads highlight the FISH signal. Error bars represent the SEM. Asterisk indicates significant difference from indicated (p < 0.05) by two-tailed Student’s t test. A minimum of 2 replicates per experiment were conducted. Scale bar represents 5 μm. Representative DNA FISH images were adjusted for figure presentation only-see materials and methods.

Because DNMT1 inhibition suppressed Dox-induced *KMT2A* alterations in cultured cells, we next assessed whether this mechanism was conserved *in vivo*. Dox treatment increases *Kmt2a* copy gains in splenic cells isolated from mice and G9a inhibition prevented these events [17]. To determine whether DNA methylation similarly contributes to this phenotype *in vivo*, mice on a mixed C57BL/6-129/Sv background were treated with DNMT1i prior to Dox administration for three consecutive days. Splenic cells were subsequently isolated, and copy number was assessed by DNA FISH (**Fig. 4E-4F**). DNMT1 inhibition significantly reduced Dox-induced copy gains *in vivo* (**Fig. 4E-4F**). Treatment with 5-azacytidine similarly suppressed Dox-induced *KMT2A* alterations (**Supplemental Fig. S4C-D**). These findings demonstrate that DNMT1-dependent DNA methylation contributes to Topo II inhibitor-induced *KMT2A* genomic alterations both *in vitro* and *in vivo*. Together with the protective effects of G9a inhibition and locus-specific TET activity, these data suggest coordinated DNA and H3K9 methylation as a potentially actionable pathway for limiting chemotherapy-induced *KMT2A* amplification and rearrangement.

## DISCUSSION

Our study identifies CTCF occupancy across the *KMT2A*-associated chromatin domain as an important determinant of local chromatin architecture and susceptibility to amplification and rearrangement. The HCT116 CTCF degron model enabled acute manipulation of CTCF, while complementary findings in RPE, primary, and in vivo systems support the broader relevance of this mechanism. We further define the coordinated balance between DNA methylation, active DNA demethylation, and H3K9 methylation as an epigenetic mechanism cooperatively regulating this state. KDM3B loss shifts the locus toward increased DNA and H3K9 methylation and reduced CTCF occupancy, whereas disruption of DNMT1-dependent methylation partially restores CTCF binding and suppresses *KMT2A* genomic alterations. The dependence of targeted DNA and H3K9 methylation-induced alterations on the complementary methylation pathway supports a model in which the combined epigenetic state, rather than either modification alone, determines *KMT2A* susceptibility.

TET-mediated methylation control has established roles in suppressing malignant hematopoietic self-renewal [30, 31]. Our findings extend this function by identifying a role for TET catalytic activity in limiting genomic alterations at a defined leukemia-associated locus. Consistent with this model, locus-targeted c-TET-dCas9 activity was sufficient to suppress *KMT2A* amplification and rearrangement under conditions that otherwise promote these events. DNMT1 inhibition also suppresses Topo II inhibitor-induced *KMT2A* alterations, linking this methylation pathway to a clinically relevant source of genotoxic stress. These findings significantly extend previous observations about epigenetic regulation of *KMT2A* amplification and rearrangement by functionally linking DNA methylation homeostasis, H3K9 methylation, and CTCF occupancy as the key coordination for control of this locus [17]. These observations illustrate a critical, unappreciated role for DNA and histone methylation balance in controlling the generation of site-specific amplification and rearrangements. Identifying the endogenous TET enzyme(s) and recruitment mechanisms that maintain this protective state will help define how active demethylation contributes to *KMT2A* protection and whether its disruption promotes these alterations in other cancer contexts.

This study coupled to a previous observation emphasize that there are initial transient *KMT2A* alterations versus inherited events [17]. Rapid degradation caused *KMT2A* copy gains and rearrangement. However, rapid CTCF recovery rescued the locus from these alterations. These collective data suggest that early events remain responsive to chromatin restoration. We speculate that this mechanism is what allows locus plasticity without creating inherited genetic events. These data also emphasize that the *KMT2A* locus is under constant regulation, which could explain why this gene is so frequently altered in cancer. The reason it accumulates preferentially in hematologic malignancies needs further consideration. Regardless of why there is cancer-type enrichment for rearrangements, this study has established that epigenetic targeting and therapeutic intervention can be used to prevent *KMT2A* alterations. Having these insights lays a foundation to better understand how this locus emerges in hematologic malignancies.

The ability of DNMT1 inhibition to suppress doxorubicin-induced *KMT2A* alterations in human cells and *in vivo* raises the possibility that DNA methylation may represent an actionable determinant to intercept therapy-associated *KMT2A* amplification and rearrangement. This opportunity is relevant to Topo II inhibitor exposure since this increases therapy-related myeloid neoplasms harboring recurrent *KMT2A* rearrangements [7, 8, 10]. DNA methyltransferase inhibitors are already used clinically in myeloid malignancies, demonstrating that DNA methylation can be pharmacologically manipulated in patients [46, 51]. In fact, multiple DNMT1 inhibitors are being leveraged in clinical trials (*e.g.*, NCT03366116, NCT04167917) [52, 53]. Understanding the best way to leverage these therapies in the clinic to control these events or other alterations will be important areas to investigate in the future. Furthermore, the protection observed with locus-targeted c-TET-dCas9 activity provides evidence that local epigenetic modulation can influence the frequency of *KMT2A* alterations and supports further investigation of more selective approaches that may limit the consequences of global epigenetic disruption. Defining treatment schedules that sustain protection against *KMT2A* alterations, while preserving chemotherapy efficacy will be important to translate these findings.

In conclusion, our findings support a conserved model across cell types and species in which the balance between DNA methylation, active DNA demethylation, and H3K9 methylation influences the CTCF-occupied chromatin state of the *KMT2A* region and the susceptibility to genomic copy gains and rearrangement. Inhibition of DNMT1 or G9a, or targeted TET activity, counteracts this state and suppresses *KMT2A* genomic alterations. Together, these findings support a role for active DNA demethylation in regulating *KMT2A* amplification and rearrangement and suggest that this locus is particularly responsive to crosstalk between DNA and H3K9 methylation states. These data also provide a collection of novel biomarkers and therapeutic targets to leverage in the regulation of *KMT2A*-induced genomic alterations. More broadly, these findings raise the possibility that the combined epigenetic state control other cancer-associated genomic regions, which will in turn influence their susceptibility to structural genomic alterations.

## Data Availability

Original Hi-C, CUT&Tag, and ChIP-sequencing data have been deposited and are publicly available in Gene Expression Omnibus (GEO) as of the date of publication. Accession numbers are listed as follows: Hi-C (GSE347563), CUTCTag (GSE346911), ChIP-Seq (GSE347282). Biomodal raw data is deposited in SRA (PRJNA1526684). This paper analyzes existing, publicly available data. These accession numbers for the datasets are as follows: HCT116 CTCF ChIP-Seq (GSE179545), HAP1 Hi-C (GSE180922), H3K9me1 ChIP-Seq (GSE210480), KDM3B ChIP-Seq (GSE71885).

## Author Contributions

J.R. Whetstine, M.A. Lineberger, Z.H. Gray, wrote the manuscript with input from authors. J.R. Whetstine conceived the study and associated concepts. J.R. Whetstine, M.A. Lineberger, Z.H. Gray, conceptualized and designed most of the experiments. M.A. Lineberger, Z.H. Gray, B.I. Ferman, E. O’Donnell, R.G. Smith, K. Schiela, C. Azadegan, M.M. Toma, E. Sumner, J. Santoro, C. Miranda, M. Ahmed, J. Wilson, C. Li, A. McKelvey, H.U. Kaniskan, W. Zhang, J. Jian, F. Puddu, M. Consugar, M. Walsh, A. Bellacosa, C. Duy, H. Lee, T. Skorski, Y. Liu, contributed to and/or conducted experiments and their interpretation within the manuscript. A. Udura, W. Lautert-Dutra, K. Chetal, and R.I. Sadreyev designed and conducted the epigenomic computational analyses. A. Udura and H. Lee conducted the TCGA analyses.

## Acknowledgements

We would like to thank the glass washing, cell culture facility, genomics resource, laboratory animal facility, and cell sorting facilities as well as the biostatistics facility, especially Michael Slifker, at Fox Chase Cancer Center for support. We would also like to thank Elena Bondarenko and Carmen Guarco for technical assistance with the manuscript. We would like to thank Drs. Capucine Van Rechem, Amy Whitaker, and Hanzhi Lou for comments on the manuscript. Work related to this study is supported by R35GM144131 (J.R. Whetstine), and NIH/NCI Cancer Center Support grant P30 CA006927 (J.R. Whetstine, T. Skorski). M.A. Lineberger is supported by TUFCCC/HC Regional Comprehensive Cancer Health Disparity Partnership NCI/NIH (U54 CA221704(5)). B.I. Ferman is supported by NIH (T32 GM142606-01). M.A. Lineberger., B.I. Ferman., E. O’Donnell, R.G. Smith acknowledge support from the C. David Allis Travel Award. R.G. Smith and W. Lautert-Dutra acknowledge support from the Temple CEI-CST Graduate Fellowship Program. Y. Liu is supported by R35GM154879 and the W.W. Smith Charitable Trust (C2407). C. Duy is supported by V Foundation (V2021-017) and the W.W. Smith Charitable Trust (C2101). R.I. Sadreyev is supported by NIH (P30 DK040561).

## SUPPLEMENTAL FIGURE LEGENDS

**Fig. S1. Rapid loss of CTCF drives *KMT2A/MLL* genomic alterations**

(A) FACS analysis of cell cycle distributions in HCT116 cells was performed following auxin-induced CTCF depletion.

(B) Hi-C analysis demonstrating altered chromatin interactions at the *KMT2A* locus following CTCF depletion for 1 hr (left) or 6 hr (middle) in one of the HCT116 cell replicates. Publicly available Hi-C data show interactions following 48 hrs of CTCF depletion in HAP1 cells (right [39]).

(C) Genome-wide comparison of CTCF CUT&Tag signal in vehicle-treated HCT116 cells versus cells treated with 5-Ph-IAA for 1 h (left) or collected 6 h after drug washout (right). Each point represents an individual CTCF peak, plotted as log₂(mean signal + 1). CTCF-A1–CTCF-T1 spanning the *KMT2A*-associated region are highlighted, with the exon 11-associated site indicated separately. Solid diagonal lines indicate equivalent signal between conditions, and dashed lines indicate ±1.5-fold differences.

(D) ChIP-qPCR demonstrating restoration of CTCF occupancy at *KMT2A* exon 11 (black) 24 hr after washout following 1 hr of 1 µM 5-Ph-IAA treatment in HCT116 cells.

Error bars represent the SEM. Asterisk indicates significant difference from indicated (p < 0.05) by two-tailed Student’s t test. A minimum of 2 replicates per experiment were conducted.

**Fig. S2. DNMT1 controls *KMT2A* genomic alterations caused by KDM3 perturbation**

(A) Schematic depicting the DNA methylation cycle and its proposed role in regulating *KMT2A* copy gains and rearrangements.

(B) CpG methylation levels (upper) and differential methylation between control and KDM3B-depleted cells (middle), shown as rolling averages across 1,500 CpGs in a second replicate experiment. The gene model is shown below. Red, green, and gray lines indicate the positions of the corresponding DNA FISH probes. Yellow shading indicates *KMT2A* exons 8-14, encompassing the BCR.RT-qPCR analysis was performed to validate gene-specific siRNA knockdowns. Samples were normalized to β-actin.

(C) RT-qPCR validation of the indicated siRNA-mediated knockdowns, *DNMT1* (left), *DNMT3A* (middle), *DNMT3B* (right). Transcript levels were normalized to β-actin.

(D) RT-qPCR analysis of *KDM3B* expression in RPE cells. Transcript levels were normalized to β-actin.

(E) RT-qPCR analysis of DNMT1 expression in RPE cells. Transcript levels were normalized to β-actin.

(F) CpG methylation levels (upper) and differential methylation between control and DNMT1-depleted cells (middle), shown as rolling averages across 1,500 CpGs in the first replicate experiment. The gene model is shown below. Red, green, and gray lines indicate the positions of the corresponding DNA FISH probes. Yellow shading indicates *KMT2A* exons 8-14, encompassing the BCR.

(G) CpG methylation levels (upper) and differential methylation between control and DNMT1-depleted cells (middle), shown as rolling averages across 1,500 CpGs in a second replicate experiment. The gene model is shown below. Red, green, and gray lines indicate the positions of the corresponding DNA FISH probes. Yellow shading indicates *KMT2A* exons 8–14, encompassing the BCR.

(H) Cell growth assay demonstrating that 100 nM DNMT1i does not alter RPE cell growth after 72 h, corresponding to Fig. 2D.

(I) FACS analysis of cell cycle distributions in RPE cells treated with 25 nM KDM3i and 100 nM DNMT1i.

(J) DNA FISH analysis of *KMT2A* copy gains and rearrangements in primary AML cells co-treated with 25 nM KDM3i and 100 nM DNMT1i. The adjacent CD3 region serves as a control.

(K) Cell growth assay in RPE cells treated with 500 nM 5-AzaC.

(L) FACS analysis of cell cycle distributions in RPE cells treated with 25 nM KDM3i and 500 nM 5-AzaC.

(M) DNA FISH analysis of *KMT2A* copy gains and rearrangements in RPE cells co-treated with 500 nM 5-AzaC and 100 nM DNMT1i. The adjacent CD3 region serves as a control.

(N) RT-qPCR validation of DNMT1 overexpression. Transcript levels were normalized to β-actin.

(O) Analysis of *DNMT3A* (left), *DNMT3B* (middle), and *DNMT3L* (right) expression in TCGA LAML samples grouped by *KMT2A* amplification status. Most samples with *KMT2A* amplification do not show increased expression of these genes. Expression is shown as RSEM values. Statistical significance was determined by Wilcoxon rank-sum test.

(P) RT-qPCR validation of *DNMT1* depletion and Halo-G9a expression in RPE cells. Relative *DNMT1* (left) and *EHMT2/G9a* (right) transcript levels are shown for the indicated conditions. Transcript levels were normalized to β-actin.

(Q) RT-qPCR validation of *G9a* depletion and Halo-DNMT1 expression in RPE cells. Relative *DNMT1* (left) and *EHMT2/G9a* (right) transcript levels are shown for the indicated conditions. Transcript levels were normalized to β-actin.

(R) DNA FISH quantification of *KMT2A* copy gains and rearrangements in control or Halo-G9a-expressing RPE cells with or without DNMT1 depletion.

(S) DNA FISH quantification of *KMT2A* copy gains and rearrangements in control or Halo-DNMT1-expressing RPE cells with or without G9a depletion.

Error bars represent the SEM. Asterisk indicates significant difference from indicated (p < 0.05) by two-tailed Student’s t test. A minimum of 2 replicates per experiment were conducted.

**Fig. S3. DNA demethylation controls *KMT2A* copy gains and rearrangements**

(A) CpG hydroxymethylation (5hmC) levels (upper) and differential hydroxymethylation between control and KDM3B-depleted cells (middle), shown as rolling averages across 1,500 CpGs in the first replicate experiment. The gene model is shown below. Red, green, and gray lines indicate the positions of the corresponding DNA FISH probes. Yellow shading indicates *KMT2A* exons 8-14, encompassing the BCR.

(B) CpG hydroxymethylation (5hmC) levels (upper) and differential hydroxymethylation between control and KDM3B-depleted cells (middle), shown as rolling averages across 1,500 CpGs in a second replicate experiment. The gene model is shown below. Red, green, and gray lines indicate the positions of the corresponding DNA FISH probes. Yellow shading indicates *KMT2A* exons 8–14, encompassing the BCR.

(C) Cell growth assay in RPE cells following 72 h of 1 uM TETi (C35) treatment.

(D) FACS analysis of cell cycle distributions in RPE cells treated with 100 nM DNMT1i and 1 uM TETi.

(E) FACS analysis of cell cycle distributions in RPE cells treated with 1 uM EHMTi and 1 uM TETi.

(F) RT-qPCR validation of *DNMT1* overexpression in RPE cells. Transcript levels were normalized to *β-actin*.

(G) DNA FISH analysis of *KMT2A* copy gains and rearrangements in RPE cells expressing c-TET-dCas9 for 24 h before induction of DNMT1 overexpression for an additional 24 h. The adjacent *CD3* region serves as a control.

Error bars represent the SEM. Asterisk indicates significant difference from indicated (p < 0.05) by two-tailed Student’s t test. A minimum of 2 replicates per experiment were conducted.

**Fig. S4. Blocking DNA methylation prevents Topoisomerase II inhibitor-induced *KMT2A/MLL* genomic alterations *in vitro* and *in vivo***

(A) RT-qPCR validation of siRNA-mediated *DNMT1* depletion. *DNMT1* transcript levels were normalized to *β-actin*.

(B) FACS analysis of cell cycle distributions in RPE cells treated with 1 pg/uL doxorubicin (Dox) and 100 nM DNMT1i.

(C) DNA FISH analysis of *Kmt2a/Mll* copy gains and the control region (*Ctrl 9*) in mouse spleen cells following 5mg/kg 5-AzaC pre-treatment before 1.5mg/kg Dox administration.

(D) Representative DNA FISH images of *Kmt2a/Mll* and the control region (*Ctrl 9*) in mouse cells following 5-AzaC pre-treatment before Dox administration.

Error bars represent the SEM. Asterisk indicates significant difference from indicated (p < 0.05) by two-tailed Student’s t test. A minimum of 2 replicates per experiment were conducted. Scale bar represents 5 μm. Representative DNA FISH images were adjusted for figure presentation only-see materials and methods.

