## Supplemental data and Methods for "DNMT1-Mediated Epigenetic Reprogramming Drives *KMT2A* Amplifications and Rearrangements"

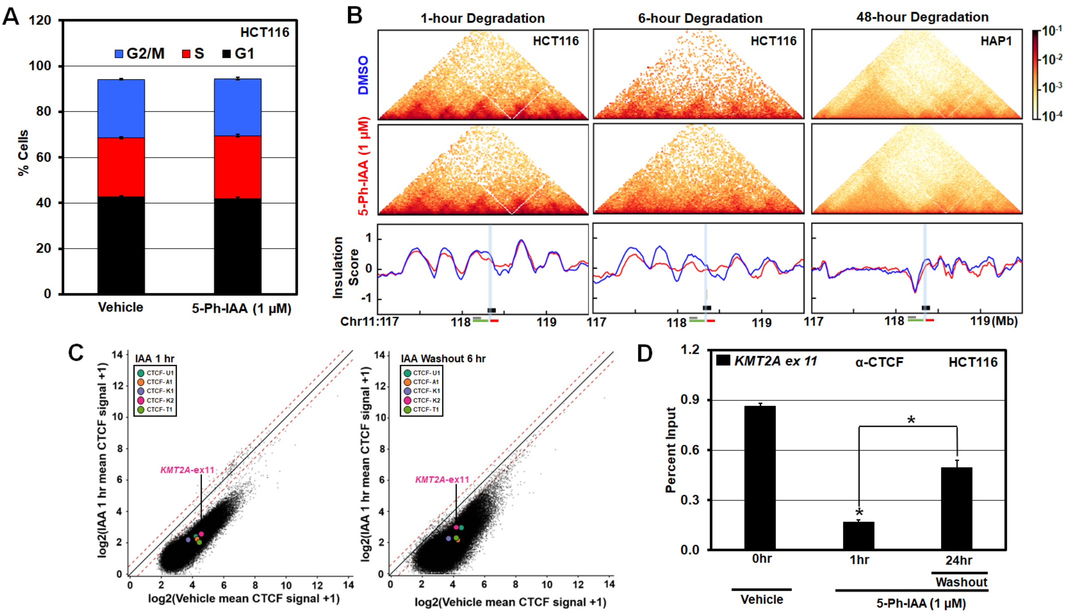
A
HCT116
G2/M
S
G1
% Cells
Vehicle
5-Ph-IAA (1 μM)
B
1-hour Degradation
6-hour Degradation
48-hour Degradation
HCT116
HAP1
DMSO
5-Ph-IAA (1 μM)
Insulation Score
Chr11:117
118
119
(Mb)
C
IAA 1 hr
IAA Washout 6 hr
CTCF-U1
CTCF-A1
CTCF-K1
CTCF-K2
CTCF-T1
KMT2A-ex11
log2(IAA 1 hr mean CTCF signal +1)
log2(Vehicle mean CTCF signal +1)
D
KMT2A ex 11
α-CTCF
HCT116
Percent Input
0hr
1hr
24hr
Washout
Vehicle
5-Ph-IAA (1 μM)

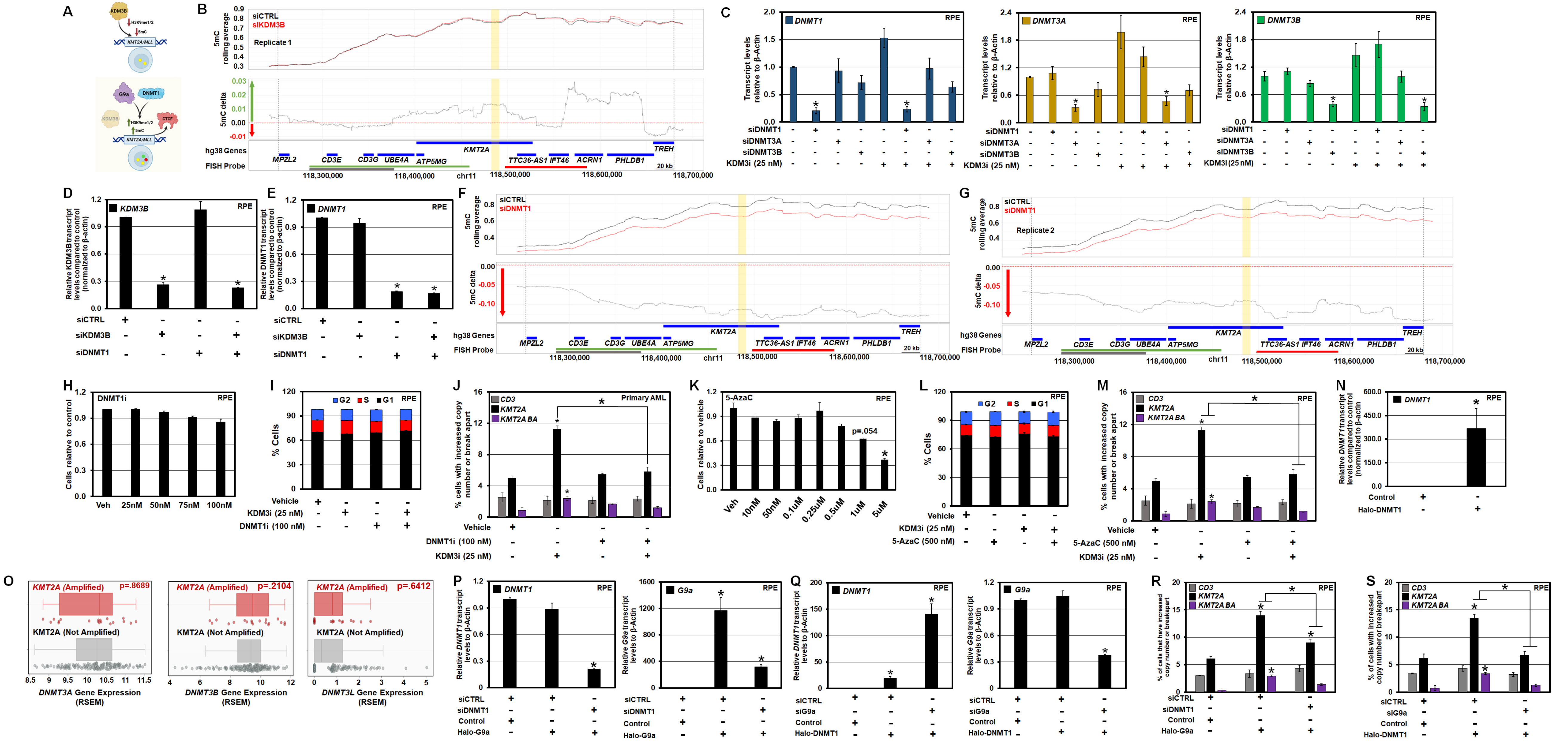
A
KDM3B
H3K9me1/2
5mC
KMT2A/MLL
G9a
DNMT1
CTCF
B
5mC rolling average
siCTRL
siKDM3B
Replicate 1
5mC delta
hg38 Genes
FISH Probe
MPZL2
CD3E
CD3G
UBE4A
ATP5MG
KMT2A
TTC36-AS1
IFT46
ACRN1
PHLDB1
TREH
20 kb
chr11
118,300,000
118,400,000
118,500,000
118,600,000
118,700,000
C
DNMT1
DNMT3A
DNMT3B
RPE
Transcript levels relative to β-Actin
siDNMT1
siDNMT3A
siDNMT3B
KDM3i (25 nM)
D
KDM3B
Relative KDM3B transcript levels compared to control (normalized to β-actin)
siCTRL
siKDM3B
siDNMT1
E
Relative DNMT1 transcript levels compared to control (normalized to β-actin)
F
siDNMT1
G
Replicate 2
H
DNMT1i
Cells relative to control
Veh
25nM
50nM
75nM
100nM
I
G2
S
G1
% Cells
Vehicle
KDM3i (25 nM)
DNMT1i (100 nM)
J
CD3
KMT2A
KMT2A BA
Primary AML
% cells with increased copy number or break apart
K
5-AzaC
Cells relative to vehicle
p=.054
10nM
50nM
0.1uM
0.25uM
0.5uM
1uM
5uM
L
5-AzaC (500 nM)
M
N
Relative DNMT1 transcript levels compared to control (normalized to β-actin)
Control
Halo-DNMT1
O
KMT2A (Amplified)
KMT2A (Not Amplified)
p=.8689
p=.2104
p=.6412
DNMT3A Gene Expression (RSEM)
DNMT3B Gene Expression (RSEM)
DNMT3L Gene Expression (RSEM)
P
G9a
Relative G9a transcript levels to β-Actin
Halo-G9a
Q
siG9a
R
% of cells that have increased copy number or breakapart
S
% of cells with increased copy number or breakapart

Supplemental Figure 3.

Lineberger et al.

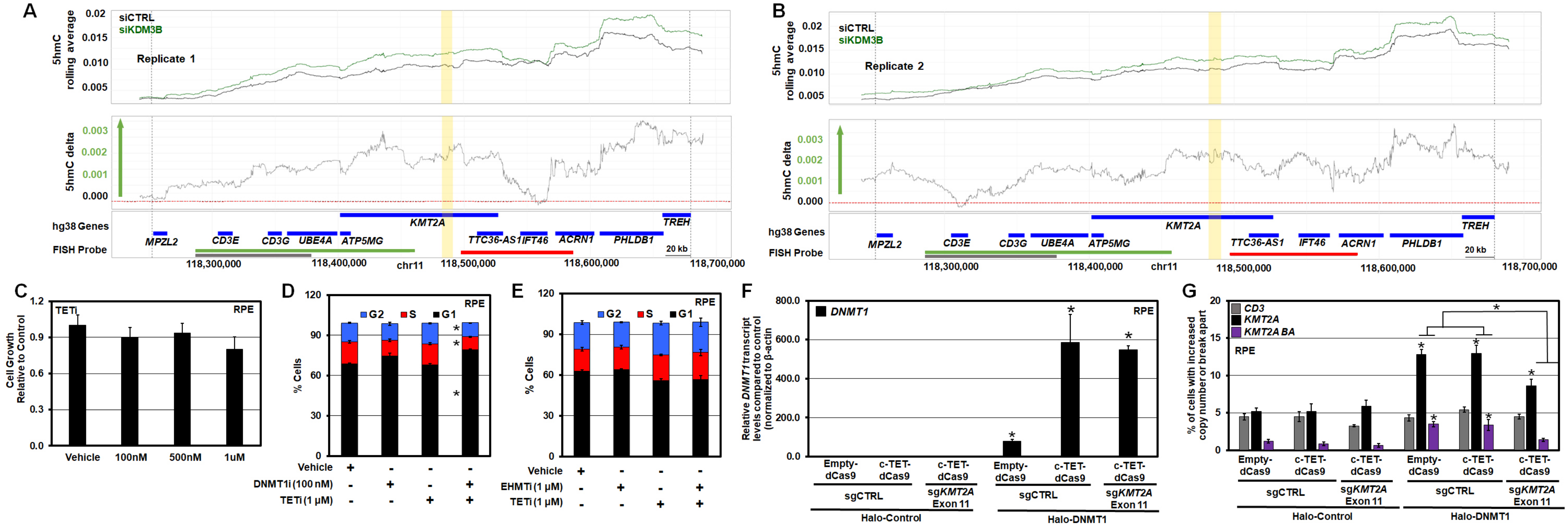

Supplemental Figure 4.

Lineberger et al.

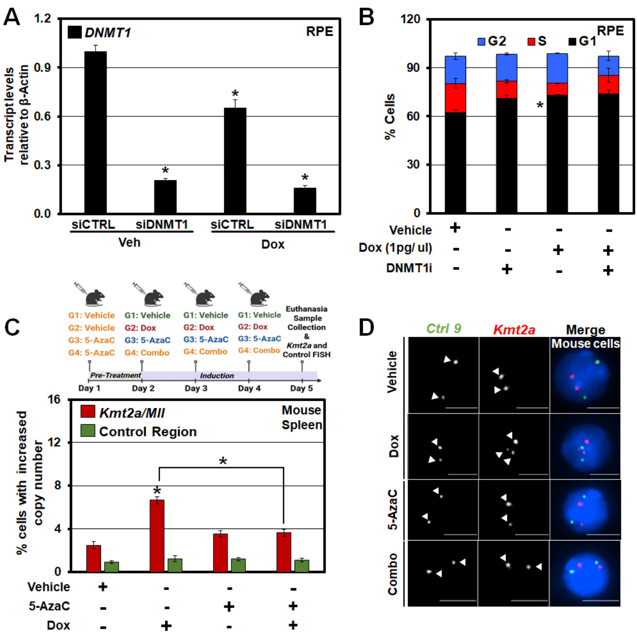

**Supplementary Table 1. siRNA Oligos**

| siRNA Name | siRNA Sequence | Unique Identifier |
|---|---|---|
| KDM3B | GGUUCACAAUCUAUACAGUtt | s28658 |
| KDM3B | UAUGCAACUGUAUAGAUUGTg | s28657 |
| DNMT1 | GCACCUCAUUUGCCGAAUAtt | s2415 |
| DNMT1 | GGAUGAGAAGAGACGUAGAtt | S2417 |
| DNMT3A | AGAUGUUCUUCGCUAAUAAtt | s200425 |
| DNMT3A | CACUGUGAAUGAUAAGCUGtt | s200427 |
| DNMT3B | GGCUAUCAGUCUUACUGCAtt | s4222 |
| DNMT3B | GCUCUUACCUUACCAUCGAtt | s4223 |
| G9a | GCUCUAACUGAACAACUAAtt | s21469 |
| G9a | CGCUGAUUUUCGAGUGUAAtt | s21470 |

**Supplementary Table 2. Oligos for qRT-PCR and qPCR**

| Primer Identifier | Primer Sequence |
|---|---|
| *KDM3B* Forward | 5'-TTATCGTGCTTCTGCTGGAA-3' |
| *KDM3B* Reverse | 5'-CATTGTGCAATGGAGACTCG-3' |
| *KDM3B* Forward | 5'-TTATCGTGCTTCTGCTGGAA-3' |
| *KDM3B* Reverse | 5'-CATTGTGCAATGGAGACTCG-3' |
| *DNMT1* Forward | 5’-AGGCGGCTCAAAGATTTGGAA-3’ |
| *DNMT1* Reverse | 5’-GCAGAAATTCGTGCAAGAGATTC-3’ |
| *DNMT3A* Forward | 5’-CCGATGCTGGGGACAAGAAT-3’ |
| *DNMT3A* Reverse | 5’-CCCGTCATCCACCAAGACAC-3’ |
| *DNMT3B* Forward | 5’-AGGGAAGACTCGATCCTCGTC-3’ |
| *DNMT3B* Reverse | 5’-GTGTGTAGCTTAGCAGACTGG-3’ |
| *G9a* Forward | 5’-TGGGAAAGGTGACCTCAGAT-3’ |
| *G9a* Reverse | 5’-GGGCAGAACCTAACTCCTCTG-3’ |
| *Actin* Forward | 5'-AGGCCAACCGCGAGAAG-3' |
| *Actin* Reverse | 5'-ACAGCCTGGATAGCAACGTACAT-3' |
| *KMT2A Ex11 CTCF site* Forward | 5’-TCTGTCACGTTTGTGGAAG-3’ |
| *KMT2A Ex11 CTCF site* Reverse | 5’-GCCCAGCTGTAGTTCTATTAC-3’ |
| *KMT2A CTCF flanking site* Forward | 5’-CAGCCAGAATCCCAGTAGA-3’ |
| *KMT2A CTCF flanking site* Reverse | 5’-CTTTCAGAGGAGGCTACAGA-3’ |

### METHODS

#### Cell Culture

Retinal pigment epithelial (RPE) cells were cultured in DMEM-high glucose (Sigma) media supplemented with 10% heat-inactivated fetal bovine serum (FBS), 100U/ml penicillin, 100µg/ml streptomycin, and 2mM L-glutamine. HL60 cells were cultured in RPMI 1640 media supplemented with 20% heat-inactivated FBS, 100U/ml penicillin, 100µg/ml streptomycin and 2mM L-glutamine. Cell line identities were authenticated by short tandem repeat analysis and Mycoplasma tested using the MycoAlert Detection Kit (Lonza, LT07-218). We appreciate the Cell Culture Facility at Fox Chase Cancer Center for their support.

Human primary patient-derived AML cells were obtained and generated as described previously [1]. Primary patient-derived AML cells were maintained on 30 Gy-irradiated OP9 feeder layer cells in complete in Iscove's modified Dulbecco's medium (IMDM; Thermo Fisher Scientific, Waltham, MA) containing 20% fetal bovine serum (Corning Premium FBS), 100 IU/ml penicillin, 100 µg/ml streptomycin, 50 µM 2-mercaptoethanol and supplemented with AML-maintaining cytokines (50 ng/ml SCF, 20 ng/ml GM-CSF, 50 ng/ml FLT3 ligand, 20 ng/ml IL-3, 20 ng/ml IL-6, and 20 ng/ml G-CSF).

HCT116 CTCF Degron cells were obtained from the Liu lab, which was kindly provided by Dr. Masato Kanemaki [2]. HCT116 degron cells were maintained in McCoys Media containing 20% fetal bovine serum (Corning Premium FBS) and 100 µg/ml streptomycin. Cell line identities were authenticated by short tandem repeat analysis and Mycoplasma tested using the MycoAlert Detection Kit (Lonza, LT07-218).

#### Transfection Procedure for RPE cells

Cells were plated in 10 cm cell culture dishes and allowed to adhere for 16-20 hrs. Cell culture medium was removed; cells were rinsed with phosphate buffered saline (PBS) and then replaced with OPTI-MEM medium (Life Technologies) prior to siRNA transfections (5nM-10nM/transfection). Transfections were changed to complete cell culture media after 4 hrs of transfection, and cells were collected 72 hrs post transfection. For **Fig. 2A**, siRNAs were transfected at the same time. Silencer select negative controls and siRNAs were purchased from Life Technologies.

Transient overexpression transfections with 1-2µg of plasmid were performed using Lipofectamine 3000 transfection reagent and P3000 reagent (Life Technologies) in OPTI-MEM medium for 4 hrs, followed by changing to complete DMEM media for 24 hrs before collection.

Transient dCas9 transfections (**Fig. 3B-E; Supplemental Fig. S3G**) with 1µg of dCas9 plasmid and 1µg sgRNA were performed using, Lipofectamine 3000 transfection reagent and P3000 reagent (Life Technologies) in OPTI-MEM medium for 4 hrs, followed by changing to complete DMEM media for 24 hrs before collection [3, 4]. Negative controls and sgRNA were purchased from IDT. **Fig. 3; Supplemental Fig. S3G** were treated with 0.75ug of dCas9 plasmid and 1ug of sgRNA.

### RNA extraction and quantitative real-time PCR

Cells were washed and collected by trypsinization after two PBS washes. Cell pellet was resuspended in Qiazol reagent (QIAGEN) for lysis and stored at -80°C before further processing. Total RNA was extracted using miRNAeasy Mini Kit (QIAGEN) with an on-column DNase digestion according to the manufacturer's instructions. RNA was quantified using NanoDrop 2000 or One (Thermo Scientific). Single strand cDNA was prepared using Super Script IV first strand synthesis kit (Invitrogen) using random hexamers. Expression levels were analyzed using FastStart Universal SYBR Green Master (ROX) (Roche) according to the manufacturer's instructions on a LightCycler 480 PCR machine (Roche) or QuantStudio 5 Real-time PCR machine (Applied Biosystems). Samples were normalized to β-actin.

### Immunoblotting

Cells were trypsinized and washed two times with PBS before resuspending in RIPA lysis buffer [50mM Tris pH 7.4, 150mM NaCl, 0.25% Sodium Deoxycholate, 1% NP40, 1mM EDTA, 10% Glycerol] freshly supplemented with Pierce Protease and Phosphatase inhibitor cocktails (ThermoFisher). Cells were lysed on ice for 15 min and stored at 80°C until further processing. Lysates were sonicated for 15 min (30sec ON and 30sec OFF cycle) at 70% amplitude in QSonica Q700 sonicator (Qsonica) followed by centrifugation at 12,000rpm for 15min. Cell lysate was transferred to a fresh tube and protein quantification was performed with Pierce BCA reagent (Thermo Scientific). Equal amounts of proteins were separated by SDS gel electrophoresis and transferred on nitrocellulose membrane (BioTrace NT, Pall Life Sciences) for at least 3 hrs at a constant current. The membranes were blocked for at least 1 hr in 5% BSA-PBST (1X PBS with 0.5% Tween-20) or 5% milk-PBST and probed over night with specific antibodies as follows at the following dilutions: anti-KDM3B (Cell Signaling) (1:1000); anti-β-Actin (Millipore) (1:10,000); anti-G9A (Sigma) (1:5000); anti-Actinin (Santacruz) (1:2000). Catalog numbers for all antibodies used in this study can be found in the Key Resource Table. Membranes were washed three times in PBST the next day, incubated with goat anti-mouse IgG peroxidase conjugated secondary antibody (170-6516, Biorad) or goat anti-rabbit peroxidase conjugated secondary antibody (A00167, GenScript) at 1:2500 in 5% milk-PBST for at least 1hr at room temperature, washed 3 times with PBST and incubated in Lumi-Light western blotting substrate (12015200001, Roche) 1min. Membranes were developed with CL-XPosure Films (34091, Thermo). The western blot images displayed in the figures have been cropped and auto-contrasted.

### Cell Cycle Analysis

Samples were washed with PBS, centrifuged at 1400rpm for 5 min, and permeabilized with 500mL PBS containing 0.5% Triton X-100 for 30 min. After this incubation, cells were washed with PBS and centrifuged at 1400rpm for 5 min. Samples were then stained with 1:100 dilutions of 1mg/mL PI solution and 0.5M EDTA with 100 mg RNase A, overnight at 4°C. Cell cycle distribution was analyzed by flow cytometry using the LSRII flow cytometry system (BD Biosciences). We are grateful to the Cell Sorting Facility at Fox Chase Cancer Center for assistance with flow cytometry.

**<u>DNA Fluorescent *In Situ* Hybridization (FISH)</u>**

The FISH protocol was performed as described previously in [5]. Briefly, cell suspensions were fixed in ice-cold methanol:glacial acetic acid (3:1) solution for a minimum of four hrs, before being centrifuged onto 8 Chamber Polystyrene vessel tissue culture treated glass slides (Falcon, Fisher Scientific) at 900rpm. The slides were air-dried and incubated in 2X SSC buffer for 2 min, followed by serial ethanol dilution (70%, 85% and 100%) incubations for 2 min each, for a total of 6 min. Air-dried slides were hybridized with probes that were diluted in appropriate buffer overnight at 37°C. The slides were washed the next day for 3 to 4 mins in appropriate wash buffers at 69°C with 0.4X SSC for Cytocell probes, Agilent Buffer1 for Agilent probes, or 0.4X SSC + 0.3% NP-40 for Empire Genomic probes followed by washing in 2X SSC with 0.05% Tween-20 (Cytocell probes), Agilent Buffer 2 (Agilent) or 2X SSC+0.1% NP-40 (Empire). The slides were incubated in 1mg/mL DAPI solution made in 1% BSA-PBS, followed by a final 1X PBS wash. After the wash, the slides were mounted with ProLong Gold antifade reagent (Invitrogen).

FISH images were acquired using an Olympus IX81 or Olympus IX83 spinning disk microscope at 40X magnification and analyzed using Slidebook 6.0 software. A minimum of 20 z-planes with 0.5µm step size was acquired for each field. For *KMT2A* break apart probe, copy gains were scored as 3 or more foci for the N terminus flanking probe (green) and C terminus flanking probe (red). Complete separation of red and green probe with no overlap was called break apart for the *KMT2A* locus with dual break apart probe. A minimum of 200 nuclei are scored for each independent experiment unless otherwise specified. Extended list of probes used are provided in the key resource table. For preparation of representative figures, DNA FISH images were uniformly adjusted for brightness and contrast to improve visualization of foci. All DNA FISH scoring and quantification were performed using the original, unprocessed images prior to any image adjustment.

**<u>Drug Treatment Conditions</u>**

Unless otherwise indicated, cells were allowed to adhere for at least 24 h before treatment. Compounds were dissolved in DMSO and added directly to complete culture medium. Corresponding vehicle-treated cells received an equivalent volume of DMSO.

**5-Ph-IAA treatment and washout.** For **Fig. 1A-D and Supplemental Fig. S1A-B**, $3.6\times10^6$ HCT116 cells were plated in 10-cm tissue-culture dishes. After 24 h, cells were treated with 1 µM 5-Ph-IAA for the indicated duration and collected. For the washout experiments in **Fig 1E-H and Supplemental Fig. S1C-D**, cells were treated with 1 µM 5-Ph-IAA for the indicated durations, washed three times with complete McCoy's 5A medium, and collected 6 h after washout, as indicated.

**KDM3 inhibition.** JDI-12 was synthesized for these studies as described previously [6]. For **Fig. 2D**, $1.5\times10^5$ RPE cells were plated in 10-cm dishes, treated with 25 nM JDI-12, and collected 24 h later. For the siRNA experiments in **Fig. 2B and Supplemental Fig. S2C**, $1.8\times10^5$ RPE cells were plated and transfected as described above. At 48 h after transfection, cells were treated with 25 nM JDI-12 and collected 24 h later.

For the 5-AzaC combination experiments in **Supplemental Fig. S2L-M**, 1.5x10$^5$ RPE cells were treated with 500 nM 5-AzaC for 48 h, followed by 25 nM JDI-12 for an additional 24 h. For the DNMT1i combination experiment in **Supplemental Fig. S2I**, 1.5x10$^5$ RPE cells were treated with 100 nM GSK3685032 for 48 h, followed by 25 nM JDI-12 for 24 h. The primary AML experiment in **Supplemental Fig. S2J** was performed using the treatment conditions described separately below.

For the epigenome-editing experiment in **Fig. 3G**, 3.0x10$^5$ RPE cells were plated and transfected as described above. Cells were subsequently treated with 25 nM JDI-12 and collected 12 h later.

**DNMT1 inhibition.** For **Fig. 3B**, 1.5x10$^5$ RPE cells were treated with 100 nM GSK3685032 for 24 h, followed by 1 µM TET inhibitor C35 for an additional 24 h. For **Fig. 4B**, 1.5x10$^5$ RPE cells were treated with 100 nM GSK3685032 for 48 h, followed by 1 pg/µL doxorubicin for 24 h. The primary AML experiment in **Fig. 4C** was performed using the conditions described separately below.

**G9a/EHMT inhibition.** For **Fig. 3C**, 1.5x10$^5$ cells were treated with 2.5 µM UNC0642 for 24 h, followed by 1 µM C35 for 12 h. For the epigenome-editing experiment in **Fig. 3E**, 2.5 µM UNC0624 was added 24 h after transient transfection, and cells were collected 24 h later.

**TET inhibition.** For the dose-response experiment in **Supplemental Fig. S3C**, 1.5x10$^5$ RPE cells were plated in 10-cm dishes and treated with the indicated concentrations of C35 and collected after 72 h. For the combination experiments in **Fig. 3B-C**, cells were pretreated with either 100 nM GSK3685032 or 2.5 µM UNC0624 before treatment with 1 µM C35, as described above.

**5-AzaC treatment.** For **Supplemental Fig. S2M**, 1.5x10$^5$ RPE cells were plated in 10-cm dishes and treated with 500 nM 5-AzaC. After 48 h, cells were treated with 25 nM JDI-12 and collected 24 h later.

**Doxorubicin treatment.** For **Fig. 4A**, 1.8x10$^5$ RPE cells were plated and transfected 24 h later. At 48 h after transfection, cells were treated with 1 pg/µl doxorubicin and collected 24 h later. For **Fig. 4B**, 1.5x10$^5$ RPE cells were treated with 100 nM GSK3685032 for 48 h, followed by 1 pg/µl doxorubicin for 24 h. For **Fig. 4D**, cells were treated with 1 pg/µl doxorubicin 24 h after transient transfection and collected 12 h later.

**Primary AML treatment.** For **Supplemental Fig. S2J**, primary AML cells were treated with vehicle (DMSO) or 100 nM GSK3685032 for 48 h, followed by vehicle (DMSO) or 25 nM JDI-12 for an additional 24 h. For **Fig. 4C**, primary AML cells were treated with vehicle (DMSO) or 100 nM GSK3685032 for 48 h, followed by vehicle or 1 pg/µL doxorubicin for 24 h before collection.

**Cleavage Under Targets and Tagmentation (CUT&Tag):**

CTCF CUT&Tag was performed in HCT116 and RPE cells using the Active Motif CUT&Tag-IT Assay Kit according to the manufacturer's instructions[7]. Following the indicated genetic or pharmacologic treatments, cells were collected, washed with ice-cold PBS, and processed using

500K cells per reaction. Nuclei were isolated and immobilized on concanavalin A-coated magnetic beads supplied with the kit.

Drosophila spike-in nuclei were added to each sample before primary-antibody incubation using the Active Motif CUT&Tag-IT Spike-In Control. Bead-bound nuclei were incubated rocking overnight at 4°C with 1ug of CTCF and 1ug of spike-in antibody. Samples were subsequently incubated with the appropriate secondary antibodies, including the goat anti-rabbit bridging antibody required for detection of the Drosophila spike-in control.

After antibody binding, nuclei were incubated with the pA-Tn5 transposase supplied with the CUT&Tag-IT kit. Following stringent washing, tagmentation was initiated using the kit-supplied tagmentation buffer. Tagmented DNA was purified and amplified using indexed primers according to the manufacturer's protocol. Amplified libraries were purified using silica purification beads per the manufacturer's protocol, quantified using Qbit, and evaluated for fragment-size distribution using TapeStation. Libraries were pooled and subjected to paired-end sequencing on an Illumina NextSeq 2000 platform using a 100-cycle sequencing kit, targeting approximately 20 million read pairs per sample. The Active Motif CUT&Tag-IT workflow and available assay formats are described by the manufacturer.

**Chromatin Immunoprecipitation Sequencing (ChIP-Seq):**

Chromatin was sonicated at 70% amplitude 15 sec on 45 sec off setting for 35 min or 45 min for CTCF ChIP. 5 μL of chromatin was RNase treated, and reverse cross-linked at least 4 hrs at 65°C in presence of proteinase K. DNA was isolated by phenol:chloroform extraction and checked on 1.3% agarose gel for a smear below 300bp. Chromatin was precleared by centrifugation at 14,000 rpm for 10 min at 4°C. Chromatin concentration was then quantified on a NanoDrop One. For each IP, 1-10μg of chromatin was immunoprecipitated with 0.2-2μg of antibody in dilution IP buffer (16.7mM Tris pH 8.0, 1.2mM EDTA pH 8.0, 167mM NaCl, 0.2% or 0.1% SDS, 0.24% Triton-X-100 or 1.84% for CTCF ChIP) at 4°C overnight. % SDS for dilution IP depended on % SDS used in Nuclear Lysis Buffer. Final concentration for IP was always 0.2% SDS. Chromatin was precleared for 2 hrs each with protein A agarose and magnetic protein A or protein G beads (Invitrogen; to match antibody isotype) rotating at 4°C before immunoprecipitation. The immunoprecipitated material was washed 2 times in dilution IP buffer, 1 time in TSE buffer (20mM Tris pH 8.0, 2mM EDTA pH8.0, 500mM NaCl, 1% Triton X-100, 0.1% SDS), 1 time in LiCl buffer (100mM Tris pH 8.0, 500mM LiCl, 1% deoxycholic acid, 1% NP40) and 2 times in TE (10mM Tris pH 8.0, 1mM EDTA pH 8.0) before elution in elution buffer (50mM NaHCO3, 140mM NaCl, 1% SDS) with RNase treatment, followed by 10μg proteinase K at 1 hr 55°C 1000 rpm. The samples were removed from beads and reverse cross-linked at 65°C for 4 hrs. Immunoprecipitated DNA was purified using either PCR purification columns (Promega) or AMPureXP beads. All the ChIPs were performed with at least two independent chromatin preparations from two independent siRNAs or two independent RPE cell lines. Antibodies used for ChIP are as follows: H3K9me1 Abcam ab8896-100. ChIP sequencing libraries were prepped using the ThruPLEX DNA-Seq kit (Takara). Libraries were paired-end sequenced (100 cycles) using a NextSeq2000 (Illumina). ChIP-qPCR was performed with 1μl of ChIP DNA.

**Biomodal duet multiomics evoC**

The duet multiomics solution evoC method (6-base sequencing [8]) was performed according to the manufacturer's instructions, using 60 ng of genomic DNA as starting material from siRNA transfected RPE cells. For primary data analysis, raw FASTQ files were processed using the Biomodal pipeline (v1.4.2) with default settings. Fastq files have been deposited in the NCBI Gene Exprression Omnibus (GEO) database and are publicly accessible. In brief, the pipeline performed adaptor trimming, resolved R1 and R2 read pairs into single-end reads retaining epigenetic information, aligned reads to the human reference genome (GRCh38), and quantified the modification state (mC, hmC, or unmodified C) at each CpG site. Methylation track plots were generating by combining counts of C, mC, and hmC across both strands at each CpG site. Additional processed filles are available upon request. The mC and hmC fractions were then calculated for sites with a minimum combined coverage of 10×. Profiles were smoothed using a rolling average spanning 1,500 consecutive CpG sites, irrespective of their genomic spacing.

**High-throughput Chromosome Conformation Capture (Hi-C)**

Hi-C experiments and analyses were performed as previously described ([9]), with minor modifications. Briefly, cells were fixed with 1% formaldehyde (Fisher Scientific, BP531-25) and disuccinimidyl glutarate (Cayman Chemical, 79642-50-5), lysed, and washed twice with cold NEBuffer 3.1 (NEB, B6003S). Nuclei were incubated with 0.1% SDS at 65 °C for 10 minutes, immediately cooled on ice, and quenched with Triton X-100 (Sigma Aldrich, 93443). Digestion was carried out by adding 400 U of DpnII (NEB, R0543M) followed by overnight incubation at 37 °C on a thermomixer. DpnII was inactivated at 65 °C for 15 minutes, and DNA overhangs were filled in with biotin-14-dATP (Active Motif, 14138) by incubation at room temperature for 4 hours. Proximity ligation was performed using 50U T4 DNA ligase (Thermo Fisher, 15224090) at 16 °C for 4 hours, followed by overnight Proteinase K (Life Technologies, 25530031) treatment at 65 °C. DNA was purified by phenol–chloroform (Life Technologies, 15593049) extraction and sonicated to an average fragment size of ~200 bp. Biotin-labeled DNA fragments were captured using streptavidin C beads (Life Technologies, 65002), and sequencing libraries were prepared by end repair and dA-tailing using the NEBNext Ultra II DNA Library Prep Kit (NEB, E6745L). Libraries were sequenced as paired-end 150-bp reads on an Illumina NovaSeq instrument.

***In vivo* Drug Treatments:**

For the *in vivo* DNMT1i combination treatment, Doxorubicin (Selleckchem) was solubilized in saline and GSK3685032 was solubilized in 10% Captisol. 16 male and 16 female mice of B6129SF1/J strain (The Jackson Laboratory 101043) were assigned into 4 groups treated with: i) vehicle (n=8, four males, four females); ii) 8 doses of GSK3685032 (twice daily, subcutaneous, 45mg/kg) (n=8, four males, four females); iii) 3 doses of Doxorubicin (daily i.v. 1.5 mg/kg) (n=8, four males, four females); iv) 8 doses of GSK3685032 with 3 doses of Doxorubicin implemented into the treatment starting day 2 (n=8, four males, four females). For the *in vivo* 5-AzaC, was solubilized in saline. 16 male and 16 female mice of B6129SF1/J strain (The Jackson Laboratory

101043) were assigned into 4 groups treated with: i) vehicle (n=8, four males, four females); ii) 4 doses of 5-AzaC (once daily, subcutaneous, 5mg/kg) (n=8, four males, four females); iii) 3 doses of Doxorubicin (daily i.v. 1.5 mg/kg) (n=8, four males, four females); iv) 4 doses of 5-AzaC with 3 doses of Doxorubicin implemented into the treatment starting day 2 (n=8, four males, four females). For both experiments, the day after the final treatment, mice were euthanized, and the cells were isolated from spleen for double blinded examination by *Kmt2a* and control FISH (*Control 9*) as previously described [10].

### QUANTIFICATION AND STATISTICAL ANALYSIS

#### DNA FISH Quantification

All pairwise comparisons were done using two-tailed Student's t-test unless otherwise stated. Significance was determined if the p value was $\leq 0.05$. All FISH experiments were carried out with at least two independent siRNAs and at least 200 nuclei per replicated were counted for all the FISH studies conducted unless otherwise stated. All FISH studies had a minimum of 2 replicates and therefore at least 400 nuclei were scored for each panel unless otherwise stated. All error bars represent the SEM.

#### Hi-C Analysis

Sequencing data were obtained in FASTQ format and trimmed to paired-end 50-bp reads using an in-house script. Trimmed reads were aligned to the human reference genome (hg19) and processed to generate multi-resolution contact matrices (mcool files) using the distiller and cooltools pipelines from Open2C [11, 12] (https://open2c.github.io/). Downstream analyses were performed using these mcool files. Insulation scores were calculated with cooltools using default parameters. Hi-C contact heatmaps and corresponding insulation profiles were generated at 20-kb resolution.

#### biomodal Analysis:

Raw FASTQ files were processed using the Biomodal pipeline (v1.4.2) with default settings. In brief, the pipeline performed adaptor trimming, resolved R1 and R2 read pairs into single-end reads retaining epigenetic information, aligned reads to the human reference genome (GRCh38), and quantified the modification state (mC, hmC, or unmodified C) at each CpG site. Differentially methylated regions (DMRs) were identified using Modality (v1.0.0). Counts of modified and unmodified cytosines at all CpG sites were aggregated within fixed-size genomic windows (1,200 bp for mC; 40 kbp for hmC). A logistic regression model was then applied to the aggregated counts, with an independent statistical test performed for each window to evaluate the null hypothesis of no differential modification.

**CUT&Tag Analysis:**

CUT&Tag sequencing reads were checked for the quality FastQC [13] and MultiQC [14]. MultiQC: summarize analysis results for multiple tools and samples in a single report. Paired-end sequences were separately aligned to the human hg19 and drosophila dm6 reference genomes using BWA-MEM [15]. Sorting and indexing was done using SAMtools [16]. *Drosophila melanogaster* spike-in reads with mapping quality ≥30 were quantified for each sample, and spike-in normalization factors were calculated relative to the sample with the lowest number of uniquely mapped dm6 reads. Human-aligned BAM files were subsequently downsampled according to these normalization factors using SAMtools to generate spike-in-normalized datasets. Deeptools bamCoverage was used to generate the normalized tracks [17]. Biological replicates were combined by arithmetic averaging, and coverage was evaluated in 200 bp bins with 600 smoothing.

Example HCT116 and RPE CTCF peaks determined from the spike-in-normalized bedGraph files using Sparse Enrichment Analysis for CUT&RUN (SEACR) (v1.3) . deepTools computeMatrix was used in reference-point mode to calculate coverage profiles within ±2 kb of the centers of selected genomic regions. The resulting matrices were visualized using deepTools plotProfile to generate signal intensity plots showing the mean normalized coverage across the selected regions as a function of distance from the 50 bp maximum peak region of the CTCF site center.

The resulting values from the matrix were transformed into $\log_2$(mean signal + 1). Scatter plots were generated using R (v 4.5.3) and ggplot2 [18] with the control condition on the x axis and the experimental condition on the y-axis. The identity line (y = x) indicated equivalent signal between conditions, while parallel dashed lines represented ±1.5-fold differences. Predefined genomic regions were highlighted as individual colored points.

**ChIP-Seq Analysis:**

ChIP-seq analysis was performed as previously described [10, 19-21]. Sequenced read quality was assessed using *FastQC* v0.12.1 and *MultiQC* v1.34 [13, 14]. Reads were processed using *fastp* v 1.3.3 for adapter trimming and low-quality read removal. Reads were then mapped to *hg19* using *bowtie2* v2.5.1 [22, 23]. Post-alignment processing, mate-fixing, coordinate sorting, duplicate marking and removal, and indexing were performed using *samtools* v1.23.1. Signal tracks of ChIP samples normalized to input were generated using deeptools v 3.5.6 (*operation* ratio and *scaleFactorsMethod* None) [17]. Count matrices used for genome-wide effect plots for target regions of the genome were generated using deeptools multiBigwigSummary.

**Schematic Diagrams:**

Schematic diagrams (**Fig. 1- 4; Supplementary Fig. S1- S4**) were created using BioRender.com

1. Duy, C., et al., *Rational Targeting of Cooperating Layers of the Epigenome Yields Enhanced Therapeutic Efficacy against AML.* Cancer Discov, 2019. **9**(7): p. 872-889.
2. Smith, R.G., et al., *Histone Acetylation Differentially Modulates CTCF-CTCF Loops and Intra-TAD Interactions.* bioRxiv, 2025.
3. O'Geen, H., et al., *dCas9-based epigenome editing suggests acquisition of histone methylation is not sufficient for target gene repression.* Nucleic Acids Res, 2017. **45**(17): p. 9901-9916.
4. Baumann, V., et al., *Targeted removal of epigenetic barriers during transcriptional reprogramming.* Nature Communications, 2019. **10**(1): p. 2119.
5. Black, J.C., et al., *KDM4A lysine demethylase induces site-specific copy gain and rereplication of regions amplified in tumors.* Cell, 2013. **154**(3): p. 541-55.
6. Xu, X., et al., *Small molecular modulators of JMJD1C preferentially inhibit growth of leukemia cells.* Int J Cancer, 2020. **146**(2): p. 400-412.
7. Kaya-Okur, H.S., et al., *CUT&Tag for efficient epigenomic profiling of small samples and single cells.* Nat Commun, 2019. **10**(1): p. 1930.
8. Füllgrabe, J., et al., *Simultaneous sequencing of genetic and epigenetic bases in DNA.* Nature Biotechnology, 2023. **41**(10): p. 1457-1464.
9. Yesbolatova, A., et al., *The auxin-inducible degron 2 technology provides sharp degradation control in yeast, mammalian cells, and mice.* Nature Communications, 2020. **11**(1): p. 5701.
10. Gray, Z.H., et al., *Epigenetic balance ensures mechanistic control of MLL amplification and rearrangement.* Cell, 2023. **186**(21): p. 4528-4545.e18.
11. Abdennur, N. and L.A. Mirny, *Cooler: scalable storage for Hi-C data and other genomically labeled arrays.* Bioinformatics, 2020. **36**(1): p. 311-316.
12. Abdennur, N., et al., *Cooltools: Enabling high-resolution Hi-C analysis in Python*. PLoS Comput Biol, 2024. **20**(5): p. e1012067.
13. Andrews, S., *FastQC: A Quality Control Tool for High Throughput Sequence Data*. 2010, Babraham Bioinformatics.
14. Ewels, P., et al., *MultiQC: summarize analysis results for multiple tools and samples in a single report.* Bioinformatics, 2016. **32**(19): p. 3047-8.
15. Li, H., *Aligning sequence reads, clone sequences and assembly contigs with BWA-MEM*, in *arXiv*. 2013.
16. Danecek, P., et al., *Twelve years of SAMtools and BCFtools.* Gigascience, 2021. **10**(2).
17. Ramírez, F., et al., *deepTools2: a next generation web server for deep-sequencing data analysis.* Nucleic Acids Res, 2016. **44**(W1): p. W160-5.
18. Wickham, H., *ggplot2: Elegant Graphics for Data Analysis*. 2 ed. 2016, New York: Springer.
19. Van Rechem, C., et al., *Collective regulation of chromatin modifications predicts replication timing during cell cycle.* Cell Reports, 2021. **37**(1): p. 109799.
20. Clarke, T.L., et al., *Histone Lysine Methylation Dynamics Control EGFR DNA Copy-Number Amplification.* Cancer discovery., 2020. **10**: p. 306-325.

21. Van Rechem, C., et al., *Lysine Demethylase KDM4A Associates with Translation Machinery and Regulates Protein Synthesis*. Cancer Discovery, 2015. **5**(3): p. 255-263.
22. Langmead, B. and S.L. Salzberg, *Fast gapped-read alignment with Bowtie 2*. Nat Methods, 2012. **9**(4): p. 357-9.
23. Chen, S., et al., *fastp: an ultra-fast all-in-one FASTQ preprocessor*. Bioinformatics, 2018. **34**(17): p. i884-i890.

**Key Resources**

| Critical commercial assays | Source | Identifier |
| --- | --- | --- |
| miRNeasy Mini Kit | Qiagen | Cat# 217004 |
| DNeasy Blood & Tissue Kit | Qiagen | Cat# 69504 |
| Superscript IV 1st Strand System | Life Technologies | Cat# 18091050 |
| CL-XPosure™ Film | Thermo Scientific | Cat# 34091 |
| Lumi-Light Western Blotting Substrate | Roche | Cat# 12015200001 |
| Pierce BCA Protein Assay | Thermo Scientific | Cat# 23223 and 23224 |
| Protein A Dynabeads | Thermo Scientific | Cat# 10002D |
| Protein G Dynabeads | Thermo Scientific | Cat# 10004D |
| CUT&Tag-IT® Express Assay Kit | Active Motif | Cat# 53177 |
| ThruPLEX® Tag-seq 96D Kit | Takara | Cat# R400585 Cat# R400586 |
| NextSeq® 1000/2000 XLEAP_SBS (100 cycles) | Illumina | Cat# 20046812 |
| evoC Biomodal Duet Multiomics | Biomodal | Cat# 6206 Cat# 4102 |
| Pierce Protease Inhibitor Tablets | Thermo Scientific | Cat# A32953 |
| Pierce Phosphatase Inhibitor Tablets | Thermo Scientific | Cat# A32957 |

| Antibodies | Source | Identifier |
| --- | --- | --- |
| Anti-JMJD1B | Invitrogen | Cat# PA5-53459 |
| Anti-beta Actin | Millipore | Cat# MAB1501; RRID:AB626633 |
| Anti-Actinin | Santa Cruz | Cat# sc-17829; RRID:AB_626633 |
| Goat anti-mouse HRP | Biorad | Cat# 170-6516; RRID: AB11125547 |
| Goat anti-rabbit HRP | GenScript | Cat# A00167 |
| Anti-CTCF, clone D31H2 | Cell Signalling | Cat# 3418 |
| Anti-H3K9me1 | Abcam | Cat# ab8896; RRID:AB_732929 |
| Anti-H3K9me2 | Abcam | Cat# ab1220; RRID:AB_449854 |
| CUT&Tag-IT® Spike-In Control, Anti-Rabbit | Active Motif | Cat# 53168 |

| Recombinant DNA | Source | Identifier |
|---|---|---|
| Halo-Tag Alone | Promega | Cat# G6591 |
| pcDNA3/Myc-DNMT1 | Addgene | Cat# 36939 |
| Empty Vector-dcas9 | Addgene | Cat# 100091 |
| DNMT3A-dCas9 | Addgene | Cat# 100090 |
| G9a-dCas9 | Addgene | Cat# 100089 |
| p-dCas9-TET1-Hygro | Addgene | Cat# 104404 |
| *MLL* breakapart | Oxford Gene Technology | Cat# LPH 013-A |
| *MLL* adjacent probe (*CD3*) | Empire Genomics | RPCI-11 215H18 |
| *Kmt2a/Chr9* Mouse | Empire Genomics | Mouse KMT2A-Chr09 |

| Deposited Data | | |
|---|---|---|
| Raw Hi-C Sequencing | This paper | GEO: GSE347563 |
| Raw ChIP-Sequencing | This paper | GEO: GSE347282 |
| Raw CUT&Tag Sequencing | This paper | GEO: GSE346911 |
| Raw DNA methylation Sequencing | This paper | PRJNA1526684 |
| HCT116 CTCF ChIP-Seq | Lee, R., et al. | GEO: GSE179540 |
| HAP1 Hi-C | Valton AL, Venev SV, Mair B, et al. | GEO: GSE180922 |
| H3K9me1 ChIP-Seq | Gray et al | GEO: GSE210480 |
| CTCF ChIP-Seq | Gray et al | GEO: GSE210480 |

| Experimental Models: Cell Lines | | |
|---|---|---|
| RPE-hTERT1 | Nick Dyson | N/A |
| HCT116 | Yu Liu | N/A |
| Primary AML | Cihan Duy | N/A |

| Experimental Models: Organisms | | |
|---|---|---|
| Mouse: C57BL/6 / 129/Sv | Jackson Labs | Strain# 101043 |

| Chemicals, Peptides and Recombinant Proteins | Source | Identifier |
|---|---|---|
| Propidium Iodide Solution | Sigma Aldrich | Cat# P4864 |
| Dimethyl sulfoxide | Millipore Sigma | D2650-100ML |
| Doxorubicin | Sigma Aldrich | ab142052 |
| Doxorubicin (Mouse study) | Selleck Chemicals | Cat# E2516 |
| 5-Azacytadine | Sigma Aldrich | Cat# A2385 |
| 5-Azacytadine (Mouse Study) | Selleck Chemicals | Cat# S1782 |
| DNMT1i (GSK3685032) | Selleck Chemicals | Cat# E1046 |
| TETi (C35) | ProbeChem | Cat# PC-72555 |

| | | |
|---|---|---|
| 5-Ph-IAA | Cayman Chemicals | Cat# 38161 |
| Dulbecco's Modified Eagles Medium – High Glucose | Sigma Aldrich | Cat# D5648 |
| McCoys 5A Modified Medium | Life Technologies | Cat# 12330031 |
| Iscove's Modified Dulbecco Medium (IMDM) | Thermofisher | Cat# 12440053 |
| Opti-Mem | Life Technologies | Cat# 31985070 |
| Trypsin (0.25%) EDTA | Life Technologies | Cat# 2520056 |
| L-Glutamine | Life Technologies | Cat# 25030-081 |
| Penicillin and Streptomycin | Life Technologies | Cat# 15140122 |
| Fetal Bovine Serum (FBS) | Gibco | Cat# 26140-079 |
| Lipofectamine 3000 | Life Technologies | Cat# L30000015 |
| EHMTi | Jian Jin | UNC0642 |
| KDM3i/JDI12 | Cynthia Myers/Jian Jin | PMID: 31271662 |

| **Software and Algorithms** | **Source** | |
|---|---|---|
| Scaffold 6.0 | 3i-intelligent imaging Innovations | https://www.intelligent-imaging.com/slidebook |
| BWA v0.7.17 | Li and Durbin, 2009 | https://bio-bwa.sourceforge.net/bwa.shtml |
| DeepTools v3.5.6 | Ramírez et al., 2016 | https://deeptools.readthedocs.io/en/develop/content/installation.html |
| FastQC v0.12.1 | Andrews, 2010 | https://www.bioinformatics.babraham.ac.uk/projects/fastqc/ |
| MultiQC v1.34 | Ewels et al., 2016 | https://github.com/multiqc/multiqc |
| fastp v 1.3.3 | Chen et al., 2018 | https://github.com/opengene/fastp |
| bowtie2 v2.5.1 | Langmead and Salzberg, 2012 | https://bowtie-bio.sourceforge.net/bowtie2/index.shtml |
| samtools v1.23.1 | Danecek et al., 2021 | https://samtools.github.io/ |
| SEACr v1.3 | Meers et al, 2019 | https://github.com/FredHutch/SEACR/ |
| bedtools v2.31.0 | Quinlan and Hall, 2010 | https://bedtools.readthedocs.io/en/stable/ |
| VennDiagram 1.8.2 | Chen and Boutros, 2011 | https://cran.r-project.org/web/packages/VennDiagram/index.html |
